# A Structure-Guided Kinase–Transcription Factor Interactome Atlas Reveals Docking Landscapes of the Kinome

**DOI:** 10.1101/2025.10.10.681672

**Authors:** Ah-Ram Kim, Kerui Huang, Jared L. Johnson, Tomer M. Yaron-Barir, Keven Wang, Lewis C. Cantley, Yanhui Hu, Norbert Perrimon

## Abstract

Protein kinases orchestrate cellular processes through phosphorylation, yet the structural basis for their specific binding partner interactions remains largely unmapped. Here, we present a structure-guided atlas of the human and *Drosophila* kinome, built by applying a new interface-aware scoring framework (iLIS) to AlphaFold-Multimer predictions. The resulting atlas recapitulates hallmark sequence preferences, confirms previously reported and functionally related protein-protein interactions, and uncovers unrecognized docking interactions. Notably, our analysis predicts a potentially widespread docking motif on homeodomain transcription factors that mediates interactions with basophilic kinases. Furthermore, we map putative allosteric interaction hotspots across the kinome and provide proof-of-concept evidence that targeting these surfaces can inhibit kinase activity. Finally, we demonstrate the physiological utility of the atlas by identifying a novel regulatory mechanism between Sgg/GSK3 and Hnf4 that controls lipid metabolism *in vivo*. This resource provides a blueprint for dissecting signaling networks and for the rational design of docking-site-specific kinase modulators.

## Introduction

The precise regulation of cellular processes hinges on protein kinases, which selectively phosphorylate their substrates to control signaling networks. While the importance of the amino acid sequence at the phosphorylation site is well-established^1–5^, a critical layer of specificity is often conferred by transient, structurally defined docking interactions at sites distant from the kinase’s catalytic site^6–8^. These interactions are challenging to capture with conventional proteomic and motif-based methods, leaving a significant gap in our ability to map signaling pathways, predict binding partner selection, and understand how kinase fidelity is precisely achieved.

Advances in deep-learning methods for protein structure prediction, such as AlphaFold^9–11^ and RosettaFold^12,13^, hold the potential to map these interactions at an unprecedented scale^14–23^. However, global confidence metrics for predicted structures are biased toward large, stable interfaces and often miss the subtle, transient docking events that are the hallmark of kinase signaling, particularly those involving intrinsically disordered regions (IDRs)^24,25^. While recent proteome-wide studies have classified IDRs into distinct “molecular grammars” based on sequence patterning^26^, the structural mechanisms by which signaling enzymes—specifically kinases—recognize and decipher these grammars remain largely unknown.

To bridge this gap and provide a structural blueprint of these interactions, we developed the integrated Local Interaction Score (iLIS). This metric identifies high-confidence interactions by combining confidence scores from both the broader interaction interface and the core physical contacts within it. To validate our framework against a challenging and biologically significant system, we applied it at a large scale to the human Ser/Thr (S/T) kinase–transcription factor (TF) interactome, modeling over 240,000 pairs. We chose to focus on kinase interactions with TFs, as they represent a critical nexus between signaling and gene expression^27,28^ and are a protein class rich in the IDRs that mediate transient interactions^24,29,30^, making them an ideal system to challenge our structure-first approach. Analysis of the resulting atlas confirmed the power of our framework, as the predicted structures successfully recapitulated established principles of kinase-partner specificity, with derived *in silico* interaction motifs aligning well with experimentally defined sequence preferences.

Next, we applied this framework to the full *Drosophila melanogaster* kinome, enabling a binding-defined kinase tree generation. This atlas reveals previously unmapped docking sites. A highlight of this analysis is the discovery of a conserved docking motif on homeodomain TFs, which reveals an unanticipated axis of structural regulation. By providing a large-scale structural view that moves beyond simple sequence motifs, this work provides a foundational resource for understanding cellular signaling and lays the groundwork for the rational design of docking site-specific kinase modulators.

## Results

### iLIS enables accurate identification of interactions involving flexible interfaces

One of the main challenges in using AlphaFold-Multimer (AFM) to predict protein-protein interactions (PPIs) lies in the limitations of global structural confidence metrics, such as the interface-predicted TM-score (ipTM)^10^, Model Confidence^10^ and pDockQ^31^. These metrics often fail to capture interactions mediated by flexible regions like short linear motifs (SLiMs) or intrinsically disordered regions (IDRs)^32^. Although AFM can model such transient complexes with high local accuracy, the presence of extensive non-interacting flexible segments can reduce overall confidence scores, pushing them below commonly accepted thresholds. To overcome this issue, we developed iLIS, a computational framework that builds upon and supersedes our previous work^33^. The iLIS score combines two measures: the Local Interaction Score (LIS; interface confidence within a PAE-defined window of ≤12 Å)^33^ and the contact-filtered LIS (cLIS; confidence restricted to residue pairs meeting both a PAE cutoff (≤12 Å) and a direct Cβ–Cβ contact ≤8 Å) (**Fig. 1A**; see **Sections 1–3 in Supplementary Text**).

**Figure 1.**
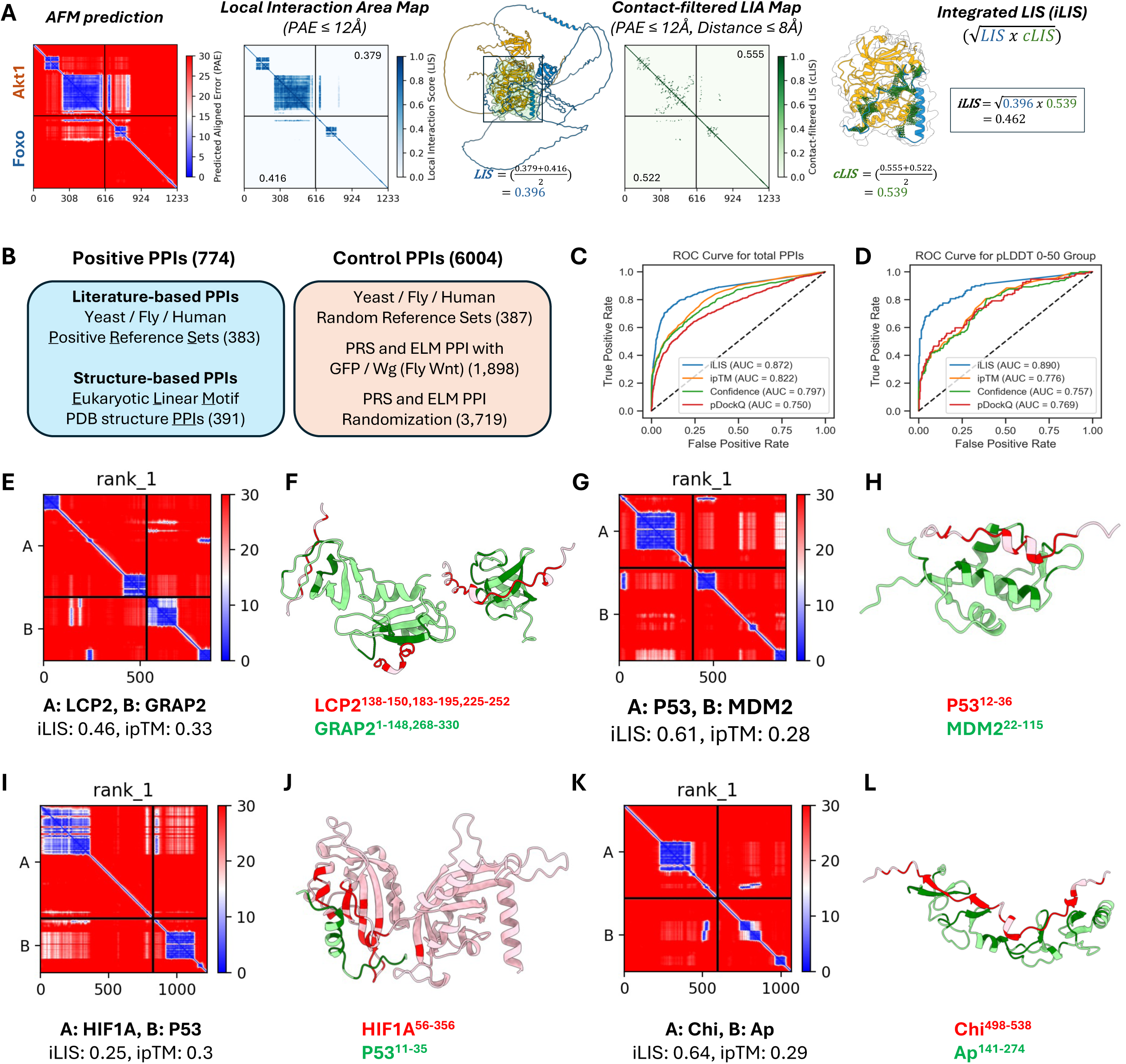
iLIS enables accurate detection of diverse protein-protein Interactions, including those involving flexible interfaces. (A) Schematic illustrating the calculation of the integrated Local Interaction Score (iLIS). AlphaFold-Multimer (AFM) predictions are first analyzed to identify the Local Interaction Area (LIA) within the complex interface, defined by a Predicted Aligned Error (PAE) cutoff (≤12 Å). The LIA is visually represented in the LIA map, where confident interaction domains in the predicted structure are highlighted by a black box. The Local Interaction Score (LIS) is then derived from the LIA by normalizing the PAE values (from the cutoff) into a confidence score between 0 and 1. LIS assesses overall interface confidence by averaging both interfaces. A contact-filtered LIA is established using both the PAE cutoff and a strict physical proximity filter (Cβ–Cβ distance ≤ 8 Å), and confident contacts are visualized by green lines in the structure. The contact-filtered LIS (cLIS) is derived by averaging the individual cLIS scores within this filtered area, focusing on direct physical contact. The geometric mean of LIS and cLIS is finally calculated to yield the robust integrated Local Interaction Score (iLIS), a metric capable of identifying both rigid and transient interactions. *Drosophila* Akt1 and Foxo interaction was used as an example here. (B) Overview of the benchmark dataset, comprising 774 positive PPIs (literature-curated and structure-based) and 6,004 control PPIs (randomized or putatively non-interacting pairs). (C, D) Receiver Operating Characteristic (ROC) curve analysis comparing the performance of iLIS to other global confidence metrics on the full benchmark dataset (C) and the disordered complexes (pLDDT 0–50) (D). (E–L) Representative examples of high iLIS-scoring interactions involving intrinsically disordered regions (IDRs) that are often missed by global metrics. Each pair includes a PAE plot (left) and a structural model (right) highlighting contacting residues. Examples shown are: (E, F) LCP2–GRAP2; (G, H) p53–MDM2; (I, J) HIF1A–p53; and (K, L) Chip–Apterous.

To evaluate the performance of iLIS, we conducted benchmarking using a diverse dataset of 774 positive and 6,004 control PPIs. This dataset was obtained from literature-curated collections used in evaluating large-scale yeast two-hybrid (Y2H) screens of the yeast, fly, and human proteomes^34–36^, as well as structure-based interactions from Eukaryotic Linear Motif (ELM) database, a resource specializing in the SLiMs that drive many transient PPIs^37^ (**Fig. 1B**; see **Supplementary Text, Sections 4–5**, for dataset details). On this benchmark, iLIS demonstrated improved performance over existing global confidence metrics like ipTM, Model Confidence, and pDockQ, achieving a ROC AUC of 0.872 versus 0.822, 0.797, and 0.750, respectively (**Fig. 1C**).

We then compared iLIS to two other recent local confidence metrics, actifpTM^38^ and ipSAE^39^. While all three local metrics are highly correlated and demonstrate similarly strong performance in distinguishing positive from control interactions (**Supplementary Fig. 1A-F**), we chose to proceed with iLIS because it holds a modest but consistent advantage in its ability to statistically distinguish between positive and control datasets (**Supplementary Fig. 1G, H**) and to recover more true positives at a 10% false positive rate (**Supplementary Fig. 1I, J**). This improved sensitivity is most apparent in challenging cases, such as highly disordered complexes (pLDDT 0–50), where local metrics are essential. iLIS maintained a high predictive accuracy (AUC = 0.890), whereas the performance of global metrics such as ipTM dropped sharply (AUC = 0.776) (**Fig. 1D**). Furthermore, iLIS demonstrated robust generalizability, outperforming ipTM on a time-split benchmark of structures deposited after AFM’s training date (**Supplementary Fig. 1K-M**). Taken together, these results highlight iLIS as a reliable and well-suited metric for identifying the transient, flexible interactions central to our study.

To visualize this feature, we highlight several bona fide interactions from our benchmark datasets that are scored highly by iLIS but poorly by ipTM (**Supplementary Fig. 2**). We defined our high-confidence thresholds at a 10% False Discovery Rate (FDR) using literature-derived Y2H reference sets, yielding cutoffs of iLIS ≥ 0.223 and ipTM ≥ 0.48. The PAE plots for these and additional examples all share a characteristic signature of localized confident interaction within complexes with uncertain global structures. Four such representative cases are visualized in **Figure 1E to 1L**. The AFM model of the LCP2–GRAP2 complex (iLIS = 0.46, ipTM = 0.33)—an interaction with a known crystal structure^40^—is correctly identified by iLIS despite a low ipTM score caused by extensive flexible regions. Similarly, iLIS successfully identifies the critical interface of the canonical p53–MDM2 interaction (iLIS = 0.61, ipTM = 0.28)^41^, where the small, structured binding motif on p53 is obscured by large disordered domains. This capability extends beyond structurally characterized pairs to known physical interactions, such as HIF1A and p53 (iLIS = 0.25, ipTM = 0.3)^42,43^, and previously mapped *Drosophila* transcription factor interactions like Chip and Apterous (iLIS = 0.64, ipTM = 0.29)^44–46^. These examples underscore the strength of iLIS in rescuing functionally critical interactions mediated by localized interfaces, a class of interactions essential to cell signaling that global metrics systematically overlook.

### Architecture and validation of the human kinase-transcription factor interactome

Having validated iLIS to identify transient, motif-driven interactions, we applied it to construct a comprehensive structural atlas of the human Serine/Threonine (S/T) kinase–transcription factor (TF) interactome. We systematically modeled over 240,000 interactions between 207 S/T kinases and 1,162 TFs. The complete set of predicted interactions, their corresponding iLIS, and the predicted residue-level interfaces for this human kinase-TF screen are provided in **Supplementary Data 1**.

The resulting interactome reveals a functionally coherent landscape where proteins with similar binding partners cluster together, correctly grouping known kinase and TF families (**Fig. 2A; Supplementary Fig. 3A**). This structure-based organization is particularly evident in our correlation analysis, which correctly clusters kinases such as the MAPKs and CDKs, and transcription factors such as the HOX family and FOXO family into their respective functional groups (**Fig. 2B, C; Supplementary Fig. 3B, C**). The network displays a hub-and-spoke architecture, with kinases like NLK and MAPK3 acting as master regulators and TFs like MYC and TP53 serving as key integrators (**Fig. 2D, E**).

**Figure 2.**
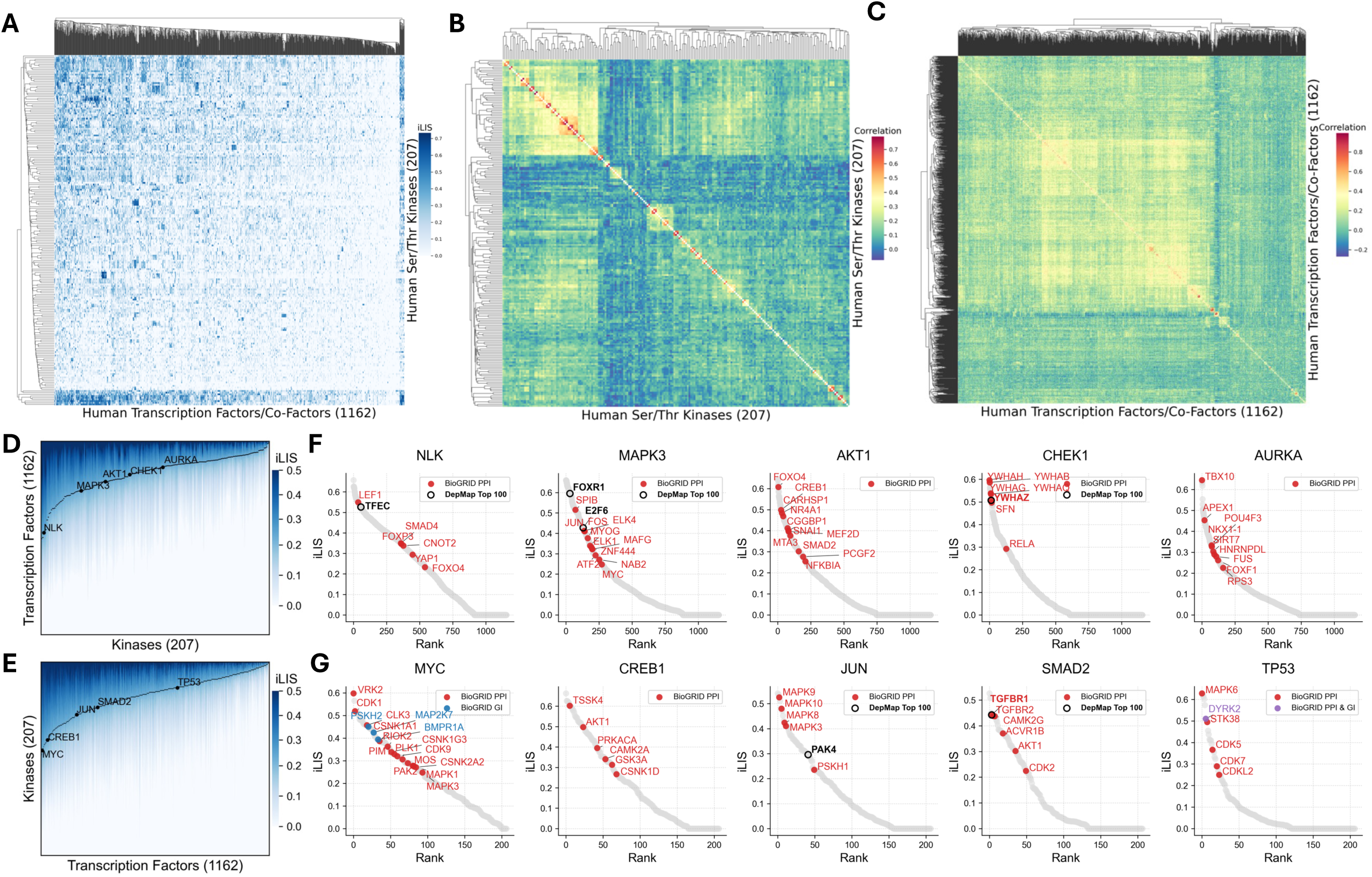
Architecture and validation of the human kinase-transcription factor interactome. (A) A comprehensive interactome of all 240,000+ predicted human S/T kinase-TF interactions. (B) Kinase correlation map by transcription factor interaction profile. (C) TF correlation map by kinase interaction profile. (D) Accumulated iLIS distribution plot of human kinases against transcription factors. Highlighted hub kinases are indicated (NLK, MAPK3, AKT1, CHEK1, AURKA). (E) Accumulated iLIS distribution plot of human TFs against kinases. Highlighted transcription factors are indicated (MYC, CREB1, JUN, SMAD2, TP53). (F) Example AFM screens of human S/T kinases against human transcription factors/co-factors. For the screens in both (F) and (G), previously reported BioGRID interactions are indicated as colored dots: Protein-Protein Interactions (PPI, red), Genetic Interactions (GI, blue), or both (purple). A black ring highlights partners that rank among the top 100 functionally co-dependent genes (ranked by absolute correlation) for that specific kinase/TF in the DepMap CRISPR screens. Highlights for external data are shown only for predictions meeting the threshold of iLIS ≥ 0.223. (G) Example AFM screens of human transcription factors against human S/T kinases.

To validate these predicted interactions, we cross-referenced the atlas against two orthogonal datasets. Our predictions recovered many known physical interactions from the BioGRID database^47^ (**Fig. 2F, G**; **Supplementary Fig. 4;** colored dots). For functional validation, we integrated our data with the Cancer Dependency Map (DepMap), which maps gene co-dependencies across diverse cancer cell lines from CRISPR screens^48^. We found that a number of our high-confidence structural predictions rank among the top 100 functionally co-dependent partners, as determined by absolute correlation score, in the DepMap dataset (25Q2), providing crucial functional support for these specific pairs and reinforcing the biological significance of the atlas. Beyond validation, the atlas serves as a powerful discovery tool: by integrating our structural predictions with known physical and genetic links, it pinpoints high-priority candidates for uncovering novel mechanisms of kinase-mediated transcriptional regulation.

### Computational motif discovery recapitulates key principles of kinase-partner specificity

From the global architecture of the kinase-TF network, we next sought to determine the structural principles that govern its interaction specificity. To do this, we derived interaction motifs from the structural interfaces for three representative kinase families (see **Supplementary Text, Section 6** for methodological details). For the basophilic kinase AKT1, our computationally derived Position Weight Matrix (PWM) (**Fig. 3A**) shows strong agreement with the known experimental motif (**Fig. 3B**)^4^, successfully recapitulating the hallmark enrichment for basic R and K residues. An interaction fingerprint highlights a specific hotspot region on the kinase (**Fig. 3C**), which corresponds to the catalytic cleft—enriched with acidic residues poised to engage the basic motifs of its partners—when visualized on the structure (**Fig. 3D**). This principle of R/K enrichment was broadly conserved across other basophilic kinases, including PKA, AKT2, PIM1, and all CDKL family members (**Supplementary Fig. 5A**).

**Figure 3.**
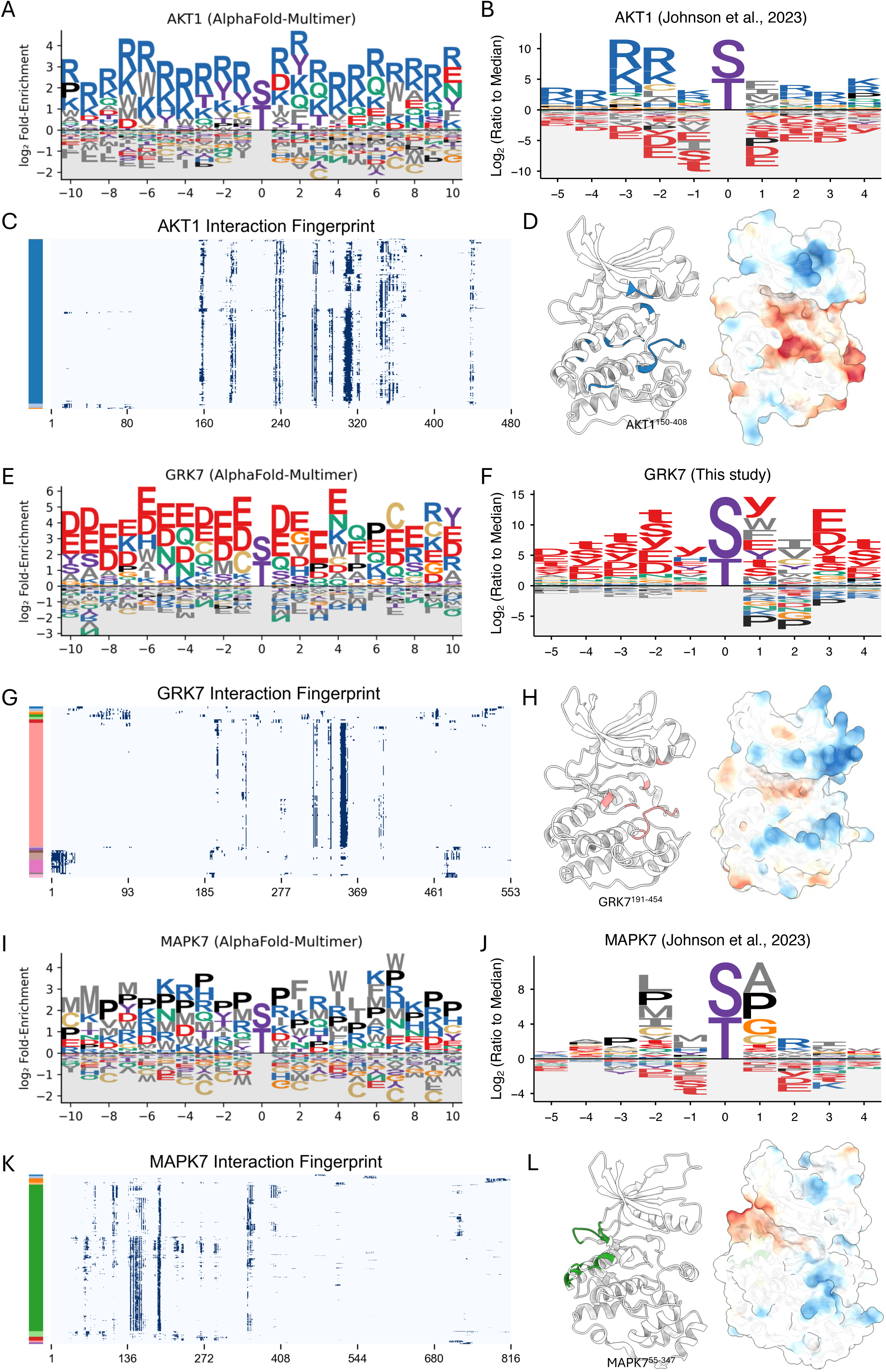
Computational motif discovery recapitulates key principles of kinase-partner specificity. (A, E, I) Computationally predicted Position Weight Matrix (PWM) for human AKT1, GRK7, and MAPK7 derived from AlphaFold-Multimer interfaces. (B, F, J) Experimentally determined PWM for human AKT1 (Johnson et al., 2023), GRK7 (This study), and MAPK7 (Johnson et al., 2023). Lowercase letters (s, t, y) denote the phosphorylated residue. (C, G, K) Interaction fingerprint of AKT1, GRK7, and MAPK7. The y-axis represents individual transcription factor partners, which are clustered based on their binding footprint on the kinase (x-axis). Color bars on the left indicate the resulting partner clusters. (D, H, L) Structural mapping of interaction hotspots. For each kinase, the left structure highlights the primary interaction hotspot, defined as residues that form contacts in at least 50% of the partners from the largest cluster in the fingerprint analysis. The right structure shows the electrostatic potential mapped to the surface (red: acidic; blue: basic). Notably, for MAPK7 (L), these hotspot residues (green) map to the conserved docking groove used for Kinase Interaction Motif (KIM) binding, a site distant from the catalytic cleft.

This principle of electrostatic complementarity is also evident for the acidophilic kinase GRK7. Our predicted PWM (**Fig. 3E**) shows a strong enrichment for the acidic residues D and E. This prediction is strongly supported by our new experimental profiling of GRK7, which confirms a clear preference for acidic phosphosites surrounding the phosphorylation site (**Fig. 3F; Supplementary Fig. 6**). The interaction fingerprint identifies specific hotspot residues (**Fig. 3G**), and the structure reveals that these hotspots form the catalytic cleft, which is lined with basic residues (**Fig. 3H**). This acidic preference was uniformly predicted across the entire GRK family and other acidophilic kinases, such as CSNK1 isoforms (CSNK1A1 and CSNK1G1) (**Supplementary Fig. 5B**).

Finally, for the proline-directed kinase MAPK7, our analysis identifies the characteristic +1 Pro preference, consistent with experimental data^4^, and a broader Pro enrichment in flanking positions (**Fig. 3I, J**). Notably, the interaction fingerprint and structure analysis for MAPK7 show that the primary hotspots are not located in the catalytic cleft but elsewhere on the kinase surface (**Fig. 3K, L**). Mapping these sites onto the structure confirms they correspond to the conserved docking groove used for Kinase Interaction Motif (KIM) binding—a mode of partner engagement separate from the active site^49–51^. This +1 Pro feature and broad Pro enrichment were also observed in other MAPK family members (**Supplementary Fig. 5C**).

While our computationally derived PWMs show general agreement with experimental data, minor deviations are observed. These may reflect the inherent limitations of our modeling, which is purely structural, considers only binary interactions (thereby excluding interactions that depend on cofactors or bridging proteins), and does not account for other cellular factors like post-translational modifications that are not captured in our models. Overall, these findings highlight the utility of a structure-based approach for large-scale motif discovery and suggest that the combination of AFM and our iLIS framework can capture key principles of kinase-partner specificity, providing a foundation for discovering novel interaction modes and regulatory networks. The full dataset containing these computationally derived PWMs for all human S/T kinases is available in **Supplementary Data 2**.

### An interactome-based kinase tree reveals binding-defined subfamilies

To systematically organize these binding principles at a systems level, we next applied our framework to the complete kinome of *Drosophila melanogaster*, performing a systems-level screen where we modeled interactions between 252 kinases (S/T, Tyr, and pseudokinases) and 841 TFs/co-factors (over 210,000 pairs). The resulting interactome provides a global view of kinome–TF connectivity (**Supplementary Fig. 7**), revealing a highly structured landscape where kinases with similar binding profiles form distinct groups. Furthermore, the network displays significant variability in connectivity, with some kinases acting as promiscuous hubs while others are highly selective, a feature also observed in our human kinome analysis. The complete prediction results for this screen, including iLIS and residue-level interfaces, are provided in **Supplementary Data 3**.

To move beyond this landscape and define binding principles, we selected the 231 kinases with sufficient data coverage for a robust clustering analysis. For this subset, we developed a pipeline using unsupervised learning to classify kinases by their binding preferences (**Fig. 4A**; see **Supplementary Text 7** for details). We first generated a quantitative ’TF-binding fingerprint’ for each kinase. These fingerprints—each one a high-dimensional vector derived from interaction hotspot data (as visualized in **Fig. 3C, G, and K**)—were then assembled into a large input matrix (231 kinases x 219,796 features) where the features represent individual TF residues.

**Figure 4.**
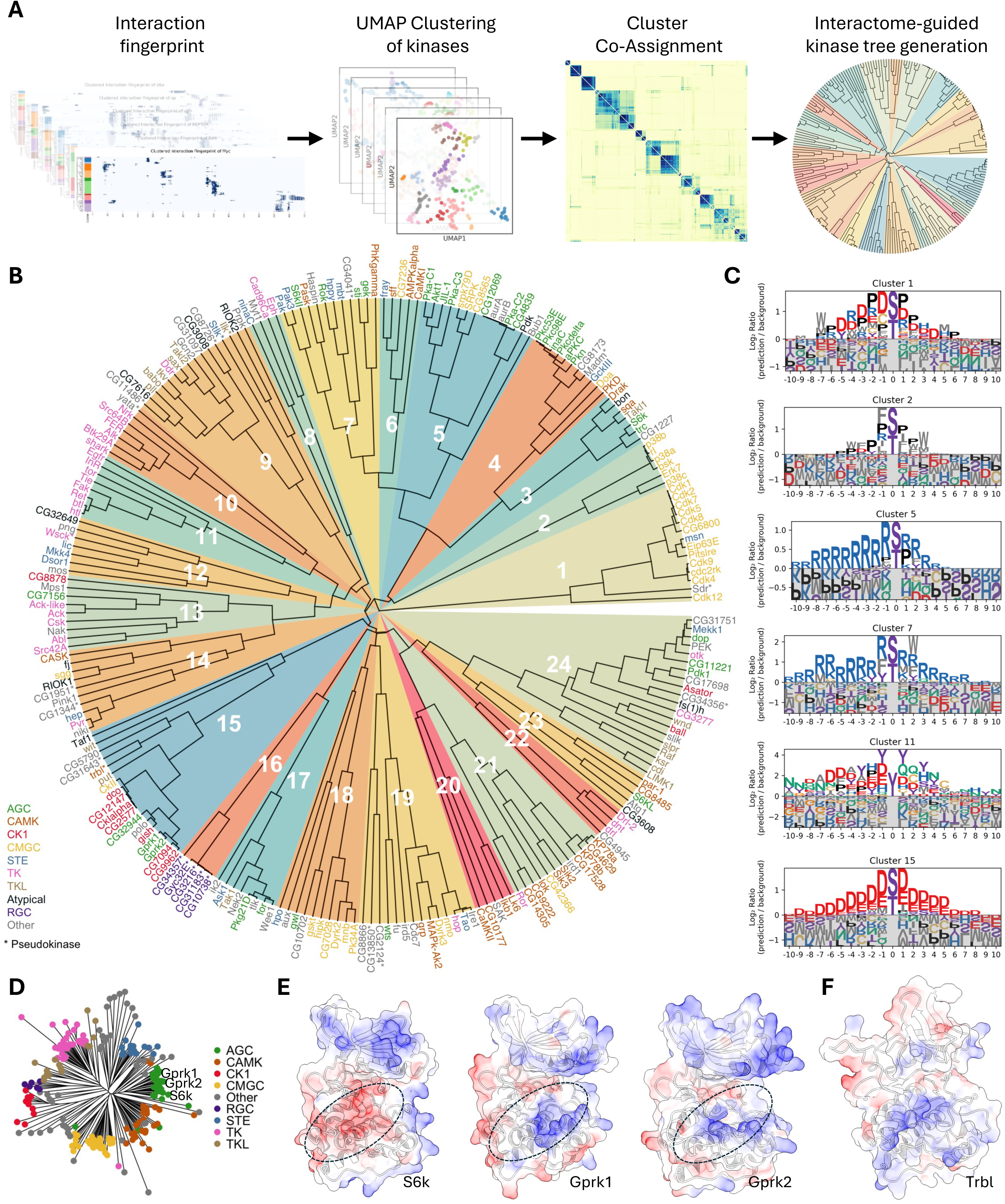
An interactome-based kinase tree reveals binding-defined subfamilies and refines functional classifications. (A) Pipeline for constructing the interactome-guided kinase tree. A “TF-binding fingerprint” for each kinase is subjected to a stability-based clustering analysis (10,800 runs) to generate a consensus matrix, which is then used for hierarchical clustering to build the final tree. (B) Binding-defined kinase tree, organizing 231 *Drosophila* kinases based on their TF-binding fingerprints. Kinases are color-coded by their sequence-based family, with pseudokinases marked by an asterisk (*). Numbered wedges indicate the binding-defined clusters. (C) Representative Position Weight Matrices (PWMs) illustrating the binding preferences for key functional clusters, such as proline-directed (e.g., Cluster 2), basophilic (e.g., Cluster 5), and acidophilic (e.g., Cluster 15) groups. The full atlas of cluster motifs is in Supplementary Figure 8. (D) Sequence homology-based kinase tree, which groups Gprk1 and Gprk2 with the basophilic AGC family. (E) Structural basis for the reclassification of Gprk1 and Gprk2. Although sequence-homologous to basophilic kinases like S6k (acidic cleft, red), their binding fingerprints place them in the acidophilic Cluster 15. This is explained by their basic catalytic clefts (circled, blue), which are suited for acidic partners. (F) The pseudokinase Tribbles (Trbl) clusters with the acidophilic group (Cluster 15). Its electrostatic surface reveals a basic pseudoactive site (circled), suggesting it retains the ability to bind acidic partners.

Because dimensionality reduction with the Uniform Manifold Approximation and Projection (UMAP) algorithm^52^ can produce variable results depending on initial conditions and hyperparameters, we employed a stability-based approach. We performed over 10,800 clustering runs, each initiated by a UMAP projection with varying parameters and bootstrapped features, and aggregated the results into a final consensus matrix. This consensus matrix robustly quantifies kinase similarity. We then used this matrix to build the final binding-defined kinase tree via hierarchical clustering. The resulting tree organizes the kinome by binding preference rather than by the sequence homology of kinase domains, helping to resolve inconsistencies in sequence-based classifications and suggesting latent functions of non-catalytic kinases (**Fig. 4B**).

This classification provides a detailed look at the kinome’s organization (**Fig. 4C**). The proline-directed kinases, for example, largely composed of the CMGC family, separate into two distinct superclusters (Clusters 1 and 2). The CDK family (e.g., Cdk1, Cdk2) forms one group that, in addition to a core proline preference, is distinguished by an enrichment for charged residues (Cluster 1). In contrast, the MAPK family (e.g., Rl, p38a) forms a second group with a clear preference for proline at canonical positions but less overall charge, instead featuring hydrophobic W and F residues (Cluster 2). Our analysis also identified a larger neighborhood of basophilic kinases, which separated into several distinct groups (Clusters 3–7). For instance, Cluster 5 (including Pka-C1, Akt1) shows a preference for a Proline at the +1 position, while Cluster 7 (a mixed group of AGC, CAMK, and STE kinases) lacks this and instead prefers hydrophobic W and F residues. Finally, tyrosine kinases formed two neighboring clusters (Clusters 10 and 11), with receptor tyrosine kinases in Cluster 11 showing a preference for tyrosine and acidic residues. Thus, the binding-defined tree not only sorts the kinome into major superfamilies but also reveals subtle, biochemically distinct binding preferences that subdivide these large classes.

### The interactome tree refines sequence-based kinase classifications

Sequence-based trees chart evolutionary history^53,54^ but can group kinases with divergent binding preferences and leave some families ambiguous (**Fig. 4D**). Clustering by interaction fingerprints offers a complementary, biochemical view, often grouping kinases from distinct sequence clades into binding-defined clusters (e.g., the mixed AGC/CAMK/STE group in Cluster 7). Our tree thus provides a framework for re-evaluating kinase family definitions and assigning previously unclassified kinases.

A compelling example of this re-evaluation is the G-protein-coupled receptor kinase (GRK) family. While sequence-based annotations place the fly GRKs, Gprk1 and Gprk2, within the predominantly basophilic AGC family (**Fig. 4D**)^55^, our interaction fingerprint analysis clusters them with canonical acidophilic kinases like the CK1 family and Polo (Cluster 15). Consistent across species, our analysis of human GRKs also revealed a clear acidophilic motif (**Fig. 3E, Supplementary Fig. 5B**) that is in strong agreement with large-scale experimental motif data^4^ and, crucially, with our new experimental data for GRK7 presented in this study (**Fig. 3F, Supplementary Fig. 6**). This finding aligns with previous functional studies showing that GRKs are, in fact, functionally acidophilic^56,57^. Our structural models of Gprk1 and Gprk2 provide a mechanistic basis for this finding, revealing a catalytic cleft lined with basic residues poised to bind acidic substrates (**Fig. 4E**).

Furthermore, because our approach is based on structurally predicted binding, it provides a novel means to characterize pseudokinases, whose binding preferences around the pseudoactive site cannot be identified by traditional phosphotransfer assays. For instance, the well-known pseudokinase Tribbles (Trbl), a member of the CAMK family, clusters with the acidophilic kinase group (Cluster 15). This suggests that Trbl retains specific binding preferences for acidophilic partners by utilizing a basic surface on its pseudoactive site (**Fig. 4F**), allowing it to function as a ‘pseudosubstrate-like decoy’ or competitive inhibitor^58–60^. By binding the same targets as active acidophilic kinases, Trbl may act as a kinase antagonist, shaping pathway output.

Finally, our tree provides a framework for classifying kinases with ambiguous assignments. An example is the Aurora kinase family (AurA/AurB). While traditional sequence homology places them in an ’Other’ group^54,55^ and a structure-based multiple sequence alignment reassigns them to the CAMK family^61^, our binding-defined tree offers a third, distinct classification. In our analysis, Aurora kinases sort into the basophilic Cluster 5, alongside kinases from the AGC and CMGC families.

In summary, this interactome-based approach provides a robust framework for classifying kinases by their binding profiles. It offers a functional view that complements traditional sequence-based trees by largely recapitulating the major kinase families while also revealing subtle subgroup preferences, as exemplified by the reclassification of the GRKs. These results suggest that this interaction fingerprinting strategy could be a broadly applicable tool for classifying other large protein families.

### A conserved docking motif links homeodomain transcription factors to diverse kinase families

While analyzing the canonical motifs from our kinase tree, we noticed that several clusters displayed unexpected preferences for residues in their flanking regions. For instance, the proline-directed Cluster 2 and the basophilic Cluster 7 both showed a distinct enrichment for hydrophobic F (phenylalanine) and W (tryptophan) residues (**Fig. 4C**). This prompted a systematic survey of our complete set of PWMs, which confirmed that motifs from multiple basophilic and MAPK clusters had recurring spikes for W, F, and Q/N in these regions (**Fig. 5A, B; Supplementary Fig. 8; Supplementary Data 4**). Observing that these modest signals often appeared linked, we hypothesized they might form a composite docking motif.

**Figure 5.**
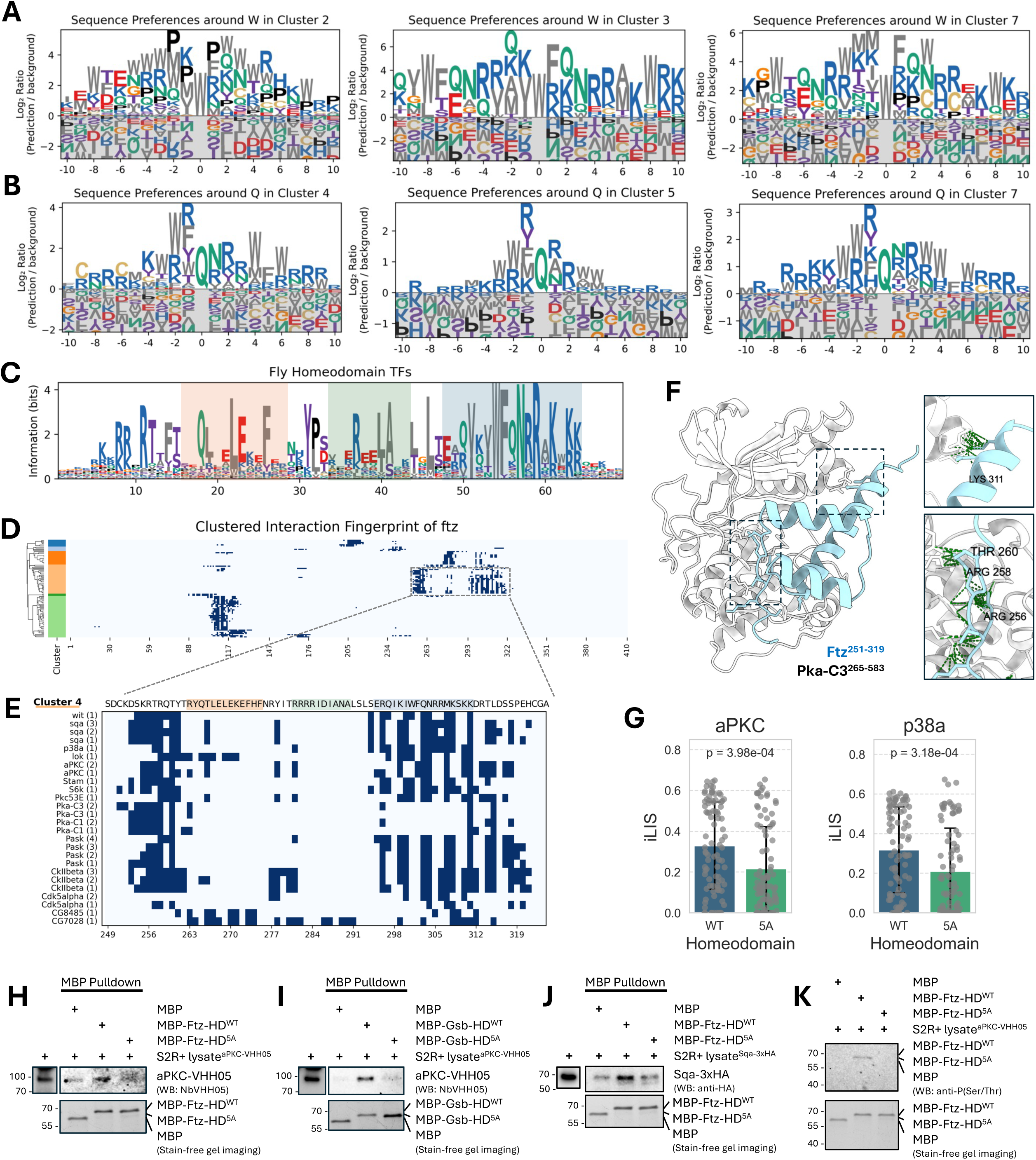
A Conserved docking motif provides a structural link between homeodomain transcription factors and kinases. (A, B) Identification of non-catalytic docking features. A survey of PWMs from multiple kinase clusters revealed recurring enrichment for flanking W, F, Q, and N residues. Representative motifs centered on W (A) and Q (B) are shown. (C) Sequence logo of a conserved composite motif identified in Homeodomain (HD) transcription factors. The logo, derived from all *Drosophila* HD proteins, highlights the consensus for the domain’s three α-helices (helix 1: pale orange; helix 2: pale green; helix 3: pale blue). (D) Clustered interaction fingerprint for the HD protein Fushi tarazu (Ftz). Kinases (y-axis) are clustered by their interaction patterns, revealing the homeodomain as a major hub. The boxed region is shown in detail in (E). (E) Focused view of the Ftz homeodomain fingerprint revealing two distinct interaction hotspots: a primary site near the N-terminus and a second corresponding to the composite motif on the third α-helix, where Lysine 311 (K311) acts as a key anchor point. Kinases are listed with their corresponding model rank. (F) Structural model of the Pka-C3–Ftz complex illustrating a two-pronged binding mechanism. The kinase’s catalytic cleft engages the N-terminal region (bottom inset), while a separate surface contacts the composite motif on the third α-helix (top inset). (G) *In silico* mutagenesis of five key basic residues to Alanine (5A) in the auxiliary motif was predicted to significantly reduce interaction strength (iLIS) with both the basophilic kinase aPKC and the MAPK p38a. (H-J) MBP pulldown assays showing basophilic kinase co-purification with wild-type homeodomains but not 5A mutants. aPKC binds Ftz-HD (H) and Gsb-HD (I); Sqa binds Ftz-HD (J). (K) *In vitro* phosphorylation assay. Wild-type Ftz-HD shows phospho-Ser/Thr signal with aPKC lysates; 5A mutant shows no phosphorylation. Stain-free gel imaging confirms equal loading.

To test this hypothesis, we performed a proteome-wide scan for this constellation of residues. The search identified a highly specific composite motif, [V/I]xxWFQNRRx[K/R]x[K/R][K/R], which maps precisely to the third α-helix of homeodomain (HD) transcription factors. A consensus sequence derived from all *Drosophila* HD TFs confirms the strong conservation of this core motif (**Fig. 5C**).

Filtering our atlas for HD-containing proteins revealed a broad regulatory network. Numerous kinases from multiple families form high-confidence interactions with dozens of HD TFs, with basophilic kinases and MAPKs being particularly enriched among the top interactors in flies (**Supplementary Fig. 9A-C**). This kinase-HD network appears to be deeply conserved, as a similar analysis of our human interactome revealed a widespread and enriched set of interactions (**Supplementary Fig. 9D-E**).

To explore the functional implications of this network, we focused on Fushi tarazu (Ftz), a key HD transcription factor required for embryonic segmentation in *Drosophila*^62–65^. The Ftz protein is known to be dynamically phosphorylated *in vivo* at numerous sites during different stages of development^66,67^. Our interaction fingerprint confirms that its HD is a major interaction hub for numerous kinases (**Fig. 5D**). A detailed view of this region reveals two primary, spatially distinct interaction hotspots (**Fig. 5E**): the first is near the N-terminus of the HD around a known PKA phosphorylation site^67^, while the second corresponds to the newly identified composite motif on the third α-helix.

Our structural model of the Pka-C3–Ftz complex provides a mechanistic basis for this dual-site interaction (**Fig. 5F**). The model shows the kinase’s catalytic cleft engaging the N-terminal phosphorylation region, while a separate surface on the kinase contacts the C-terminal composite motif on the third α-helix. This suggests a model where the N-terminal region serves as the primary catalytic site, while the conserved basic patch within the newly identified composite motif is predicted to act as an auxiliary docking site to enhance binding affinity and specificity. This two-pronged binding mode appears to be a general feature, as similar dual interaction hotspots were observed for numerous other HD TFs in both fly and human (**Supplementary Fig. 10**).

To test the functional importance of the conserved basic patch within the C-terminal composite motif, we first performed *in silico* mutagenesis of its five key basic residues to Alanine (5A). We screened these 5A mutants from all fly HD proteins against aPKC (a basophilic kinase) and p38a (a MAPK), the top-enriched HD-interacting kinases from their respective families. The models predicted that the 5A mutation would broadly disrupt these interactions, as reflected by a significant reduction in iLIS scores (**Fig. 5G**). Since mutating these basic residues *in vivo* would create confounding effects by disrupting DNA binding, we performed biochemical validation using purified, DNA-free MBP-tagged homeodomain fragments from two fly HD proteins, Ftz and Gooseberry (Gsb). First, we carried out pulldown assays to test PPIs. Wild-type HD fragments from both Ftz and Gsb specifically co-purified basophilic kinases (aPKC and Sqa) from transfected S2R+ cell lysates, whereas the 5A mutants failed to support detectable kinase recruitment above background levels (**Fig. 5H-J**). To determine whether this binding enabled phosphorylation, we incubated the fragments with aPKC-containing lysates. We observed a phospho-Ser/Thr antibody signal for the wild-type Ftz fragment, but this phosphorylation was absent in the 5A mutant (**Fig. 5K**). Together, these results indicate that the basic patch is required for kinase recruitment and subsequent substrate phosphorylation across HD-kinase pairs, establishing it as a functional docking interface.

These findings reveal a potential mechanism for integrating signaling inputs with transcriptional control. While the third α-helix is essential for DNA binding^68–72^ and serves as an interface for PPIs with other cofactors^73–75^, our data add a critical new dimension to its function: it also acts as a docking platform for basophilic kinases. Our models and *in silico* mutagenesis indicate that the conserved basic patch within this composite motif stabilizes kinase, and our biochemical experiments confirm this interaction is necessary for substrate phosphorylation. Together, these findings establish a model where kinases dock onto a multifunctional surface—one that mediates DNA binding, cofactor interactions, and kinase recruitment—positioning them to phosphorylate their targets and regulate the transcriptional activity of the HD protein. Mutations in this critical region are therefore likely to have pleiotropic effects by disrupting both kinase docking and DNA recognition, which may explain the high prevalence of pathogenic variants at this site^72,76^.

### The kinome atlas systematically identifies inhibitable allosteric hotspots

A central challenge in drug development is designing selective kinase inhibitors, as the ATP-binding catalytic site is highly conserved^77,78^. Targeting unique, non-catalytic (allosteric) surfaces offers a promising route to achieving specificity^79–81^, but a systematic method for identifying such sites has remained elusive.

Our atlas addresses this challenge by systematically mapping interaction hotspots across the entire kinase surface, revealing two principal strategies for partner recognition that are distinguished by their binding location. The first and most common strategy utilizes the catalytic cleft, which serves as the primary, electrostatically potent interface for kinases that prefer highly charged partners, such as the basophilic and acidophilic families (**Fig. 6A, B; Supplementary Fig. 11A, B**). In a second, contrasting strategy, kinases rely on auxiliary (allosteric) docking sites for partner engagement. This mode is employed by kinases that favor less-charged motifs, like the MAPK families (**Fig. 6C**; **Supplementary Fig. 11C**), but is also a noteworthy solution used by certain acidophilic kinases like CSNK2A1 and PLK1, demonstrating that diverse structural solutions for partner recognition exist (**Supplementary Fig. 11D**). This systematic classification of binding modes provides a powerful framework for identifying functionally critical—and potentially inhibitable—allosteric surfaces across the kinome.

**Figure 6.**
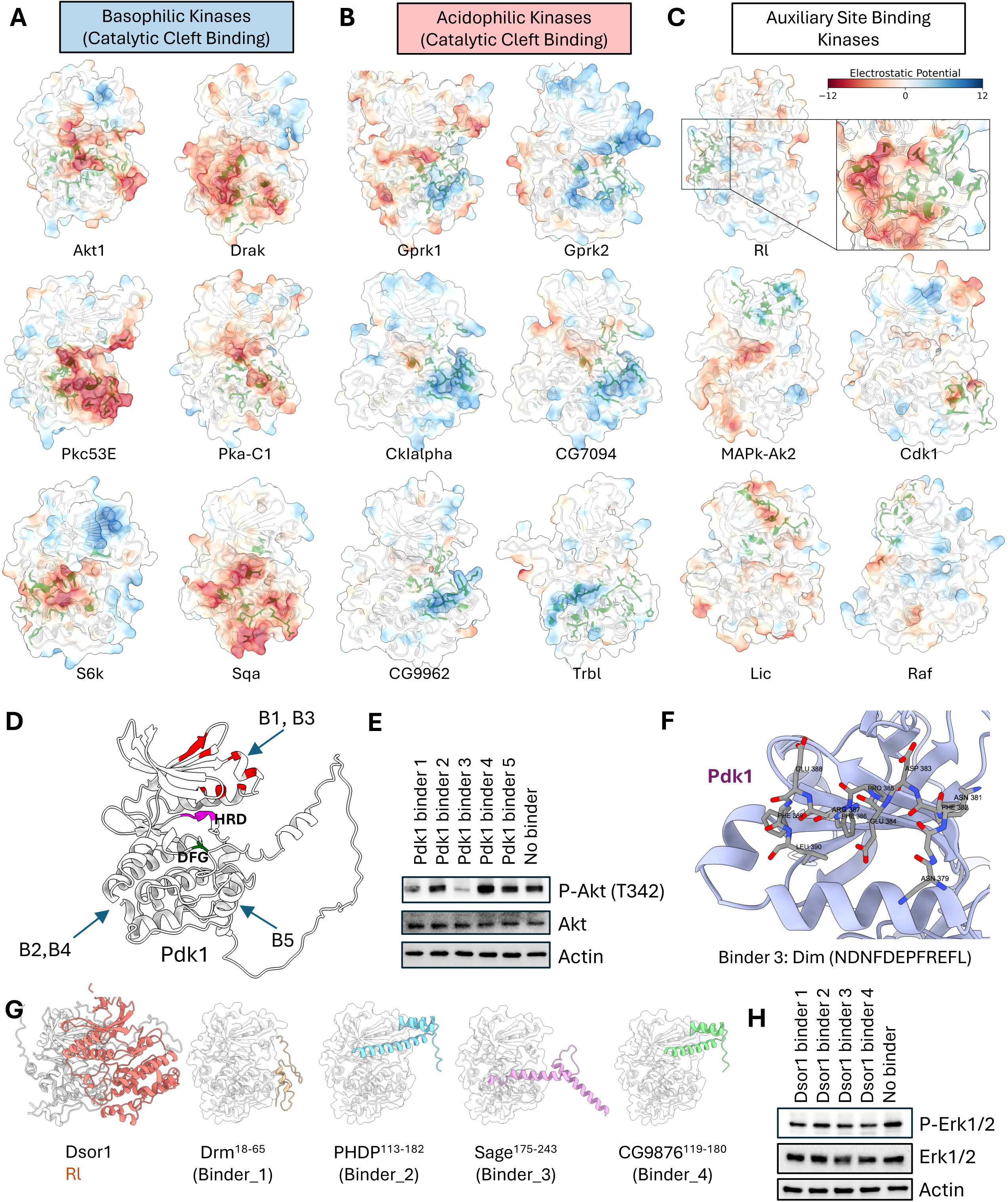
The kinome atlas identifies functionally relevant and inhibitable docking hotspots. (A–C) Principal modes of kinase partner recognition with electrostatic potential mapped to the kinase surface (red: acidic; blue: basic). (A) Basophilic kinases using the catalytic cleft. (B) Acidophilic kinases using the catalytic cleft. (C) Auxiliary site-binding kinases using non-catalytic (allosteric) surfaces. (D–F) Validation of a predicted allosteric hotspot on Pdk1. (D) Model of Pdk1 with interaction hotspots highlighted in red and predicted binding sites for five TF-derived fragments shown. Binders 1 & 3 target the allosteric PIF pocket. (E) Western blot showing that only Binders 1 & 3 inhibit downstream phosphorylation of Akt (at T342) in S2R+ cells. (F) Structural model of Binder 3 docked within the Pdk1 PIF pocket. (G–H) Validation of a predicted MAPK-docking site on Dsor1. (G) Models of the MAP2K Dsor1 and four TF-derived fragments predicted to inhibit Rl binding. (H) Western blot showing reduced phosphorylation of Rl (Erk1/2) in S2R+ cells upon expression of all four binder fragments.

To confirm that these predicted hotspots are functionally relevant and represent potential inhibitory targets, we experimentally validated two distinct sites. First, we focused on Pdk1. Its N-lobe contains the well-characterized PIF pocket, a docking site that is required for the phosphorylation and activation of key downstream substrates, including Akt1. Prior studies have shown that synthetic peptides like PIFtide can competitively inhibit this site, blocking downstream signaling and establishing it as a bona fide inhibitory target^82,83^. Consistent with this, our atlas identified the PIF pocket as the primary interaction hotspot on Pdk1 and predicted that fragments from several TFs would bind there. To test these predictions, we selected five TF-derived fragments: two targeting the PIF pocket hotspot, and three targeting other surfaces as controls (**Fig. 6D**). When expressed as BFP fusion proteins in *Drosophila* S2R+ cells, only the two fragments targeting the PIF pocket (Binders 1 and 3) effectively inhibited Pdk1’s downstream phosphorylation of Akt1, whereas the control fragments had no effect (**Fig. 6E**). Our structural models predict that these inhibitory peptides dock directly within the PIF pocket, providing a clear mechanistic basis for their competitive effect (**Fig. 6F**).

Second, we investigated a predicted hotspot on the MAP2K, Downstream of raf1 (Dsor1), required for the activation of its downstream MAPK, Rl. Among Dsor1-interacting TFs, we selected four high iLIS-scoring TF fragments. Our models predicted that these fragments would bind to the Dsor1 surface that engages Rl, suggesting they could act as competitive inhibitors (**Fig. 6G**). To test this, we expressed the fragments as BFP fusions in S2R+ cells. As predicted, all four fragments reduced phosphorylation of the downstream MAPK, Rl (Erk) (**Fig. 6H**). Notably, two of these inhibitory fragments were derived from homeodomain TFs (PHDP and CG9876), suggesting that in addition to being kinase targets, homeodomains may possibly, in certain contexts, also function as allosteric modulators of kinase activity.

Collectively, these results, where computational predictions for two distinct kinase families were supported by functional experiments, demonstrate the use of our atlas for the systematic discovery of functional, non-catalytic sites. We provide a comprehensive catalog of all interaction hotspots—encompassing both catalytic and putative allosteric surfaces—across the *Drosophila* and human kinomes (**Supplementary Data 5**), which can serve as a structural guide for developing a new generation of highly selective, allosteric kinase modulators.

### The kinome**–**TF atlas pinpoints regulators of Hnf4 and lipid homeostasis

To test if our kinome–TF atlas could uncover physiologically relevant connections, we sought to identify novel regulators of *Drosophila* Hepatocyte Nuclear Factor 4 (Hnf4), a key transcription factor controlling lipid metabolism^84–86^. We implemented a two-step filtering strategy, first identifying kinases predicted to interact with Hnf4 with high confidence (iLIS ≥ 0.223) and then intersecting this list with transcriptomic data from oenocytes to narrow down to 18 co-expressed kinases for functional validation (**Fig. 7A**). Oenocytes are a principal metabolic tissue in *Drosophila*, serving a role analogous to the mammalian liver. An adult oenocyte-specific RNAi screen of these candidates in endogenously NanoTagged Hnf4 flies (Hnf4-127D01)^87,88^ revealed that knockdown of 12 kinases (a 66% hit rate) altered Hnf4 protein abundance, underscoring the predictive power of our structure-guided approach (**Fig. 7B, C; Supplementary Fig. 12**).

**Figure 7.**
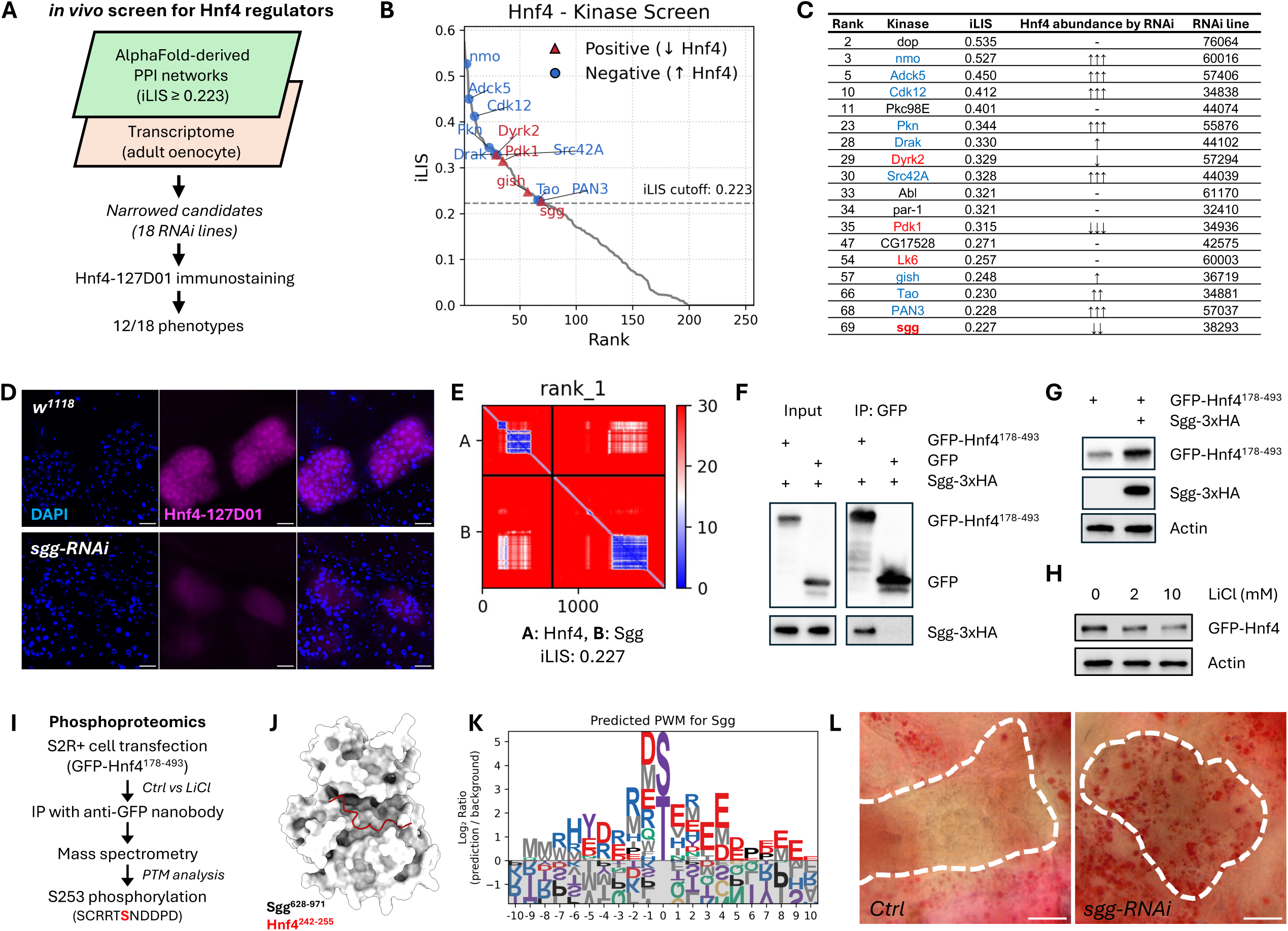
The kinome atlas identifies Sgg/GSK3 as a key regulator of Hnf4 stability and lipid homeostasis. (A) Schematic of the two-step *in vivo* screen to identify Hnf4 regulators. (B) Predicted iLIS scores for the 18 candidates from the *in silico* screen. Points are color-coded by their functional effect in the subsequent *in vivo* RNAi screen, and only candidates that produced a phenotype are labeled. (C) Summary of the *in vivo* RNAi screen results for the 18 candidates. *PromE-GAL4; tub-GAL80^ts^* was used for oenocyte-specific RNAi. (D) Immunostaining of Hnf4 (magenta) in oenocytes from control (*w¹¹¹*U) and *sgg*-RNAi adults. DAPI (blue) stains nuclei. Scale bar, 20 µm. (E) Predicted Aligned Error (PAE) map of the Sgg-Hnf4 prediction. (F) Co-immunoprecipitation (co-IP) of Sgg-3xHA with GFP-Hnf4[178-493] from S2R+ cell lysates. (G) Western blot showing stabilization of the GFP-Hnf4[178-493] fragment upon Sgg-3xHA co-expression in S2R+ cells. (H) Western blot showing a reduced level of full-length GFP-Hnf4 in S2R+ cells after treatment with the GSK3 inhibitor LiCl. (I) Schematic of the phosphoproteomics workflow identifying Serine 253 (S253) as an Sgg-dependent phosphorylation site on Hnf4. (J) Structural model of the predicted Sgg-Hnf4 interface, highlighting the location of the S253-containing peptide (red). (K) Computationally derived PWM for Sgg from the fly kinome atlas. (L) Oil Red O staining of adult oenocytes, showing increased lipid accumulation (red) in *sgg*-RNAi compared to control. Knockdown was induced using the *PromE-GeneSwitch-Gal4* system; *sgg*-RNAi flies were treated with RU486 while control flies were not.

Among the positive regulators identified in this screen was Shaggy (Sgg). We prioritized Sgg for in-depth validation because its mammalian ortholog, GSK3, is a key modulator of glycogen and lipid metabolism and is known to promote inflammation in pathologies like fatty liver disease^89,90^, making it a prime candidate for a conserved regulatory role. Confirming its importance, oenocyte-specific depletion of *sgg* led to a near-complete loss of Hnf4 protein *in vivo* (**Fig. 7D**), indicating Sgg as a critical regulator of Hnf4.

Guided by the structural prediction (**Fig. 7E**), we proceeded to dissect the mechanism of this interaction. We first validated a physical interaction in S2R+ cells, showing that Sgg co-immunoprecipitated with a GFP-tagged Hnf4 fragment (Hnf4[178-493]) that contained the local interaction area with Sgg (**Fig. 7F**). Next, we confirmed that Sgg’s kinase activity is required for Hnf4 regulation. While Sgg overexpression stabilized the Hnf4[178-493] fragment (**Fig. 7G**), inhibiting GSK3 with lithium chloride (LiCl) reduced full-length Hnf4 protein levels (**Fig. 7H**).

To identify the phosphorylation site, we performed phosphoproteomics on the immunoprecipitated GFP-Hnf4[178-493] fragment. This analysis identified Serine 253 (S253) as a site whose phosphorylation is dependent on Sgg/GSK3 activity, as its signal was rendered undetectable by LiCl treatment (**Fig. 7I**). Typically, GSK3 activity is priming-dependent, requiring a prior phosphorylation event by a different ‘priming kinase’ on a serine or threonine located at the +4 position relative to the target site, creating the canonical S/T-x-x-x-pS/pT docking motif for Sgg/GSK3^91–93^. The sequence surrounding S253 does not conform to this canonical motif. However, priming-independent mechanisms, in which GSK3 can phosphorylate a substrate without a priming event, have been documented for other substrates such as Myc, Jun, and Tau^94,95,93^_._

Consistent with such a mechanism, the sequence C-terminal to S253 features an acidic patch (NDDPD). Our structural model predicts that this Hnf4 peptide docks near the catalytic site of Sgg (**Fig. 7J**). This principle is corroborated by our computationally derived PWM for Sgg, which reveals a strong general preference for acidic residues downstream of the phospho-acceptor (**Fig. 7K**). These converging lines of evidence suggest a mechanism wherein Sgg directly phosphorylates Hnf4 at S253 to control its stability.

Finally, to determine the physiological consequence of this Sgg–Hnf4 regulatory axis, we depleted *sgg* specifically in oenocytes. This manipulation induced marked lipid accumulation, a steatosis-like phenotype that phenocopies *Hnf4* loss (**Fig. 7L**)^84,85^. Together, these data validate Sgg as a critical kinase that stabilizes Hnf4 via a priming-mimetic phosphorylation at S253, and demonstrates the use of our atlas for discovering novel kinases that control organismal metabolism *in vivo*.

## Discussion

Here, we present the first systematic, structure-guided kinase–TF interactome atlas, contributing to our understanding of a longstanding challenge in cell signaling. By pairing AlphaFold-Multimer with our iLIS framework, we address a key limitation of existing metrics in recognizing interactions mediated by intrinsically disordered regions. The robustness of the resulting atlas is validated against multiple benchmarks, where it recovers known physical interactions from BioGRID^47^ and corresponds with functional co-dependencies from DepMap^48^. This work therefore provides a map of the kinome–TF interactome that captures the flexible, transient interactions underpinning signaling specificity.

A key advance of this atlas is its ability to move beyond binary interaction mapping to reveal how and where interactions occur, resolving specific interfaces that have remained elusive in large-scale screens. The atlas provides residue-level structural insight into partner recognition, exemplified by our binding-defined classification of the kinome, the discovery of a putative conserved docking mechanism for homeodomain TFs, and the systematic mapping of allosteric hotspots. The homeodomain–kinase analysis, supported by extensive computational modeling and *in silico* mutagenesis, proposes a detailed hypothesis for how these developmental regulators are engaged by upstream pathways. We provide experimental support for this mechanism through an *in vitro* phosphorylation assay, demonstrating that the basic patch in the third α-helix is required for proper phosphorylation by basophilic kinases.

We demonstrate the atlas’s power as a hypothesis-generation approach by validating *in vivo* a high-confidence predicted interaction between Sgg and Hnf4. Our platform successfully guided the identification of a specific, Sgg-dependent phosphorylation site at Serine 253. The biochemical features of this site were consistent with the binding preferences for Sgg derived from our atlas, elucidating a non-canonical, priming-mimetic phosphorylation mechanism that controls lipid metabolism *in vivo*. This mechanism is consistent with studies of other GSK3 substrates, where acidic residues can substitute for a priming phosphate, although often with reduced kinetic efficiency^96^. The physical interaction we observed between Sgg and Hnf4 may compensate for this potential inefficiency through induced proximity, increasing the effective local concentration of enzyme and substrate to drive physiologically relevant phosphorylation *in vivo*.

This structural atlas lays the groundwork for advances in drug development. While achieving selectivity against the conserved ATP-binding pocket is a persistent challenge^97–99^, our atlas provides a detailed resource for targeting unique, non-catalytic (allosteric) surfaces. With recent advances in designing bespoke protein, peptide, and small-molecule binders for specific target proteins^100–109^, the primary hurdle is shifting from how to target a surface to which surface provides the desired therapeutic effect. Our work provides a catalog of these surfaces across human and fly kinases, paving the way for the development of selective, non-ATP-site kinase modulators.

Our approach has some inherent limitations. Its accuracy is fundamentally dependent on the underlying AlphaFold-Multimer predictions, and our models analyze binary interactions in isolation, without crucial cellular context like co-factors or subcellular compartmentalization. A key limitation is the absence of post-translational modifications in our models, such as the priming phosphorylation events required by kinases like GSK3. While this prevents the detection of strictly priming-dependent interactions, it creates a productive bias that enriches for alternative, priming-mimetic recognition mechanisms. As demonstrated with the Sgg–Hnf4 interaction, our platform successfully identifies cases where the kinase recognizes a patch of acidic residues that functionally mimics a priming phosphate, positioning the atlas as a unique resource for exploring these non-canonical modes of kinase regulation. Future integration with large-scale phosphoproteomic datasets will be a powerful next step to systematically distinguish between priming-mimetic docking and true enzymatic substrates.

Beyond the limitations of the structural models, it is critical to distinguish a predicted docking event from a functional enzymatic one, as high-confidence binding does not guarantee phosphotransfer. Well-characterized pseudosubstrates, such as the Protein Kinase Inhibitor (PKI) peptide, exploit this principle by binding a kinase’s active site without being phosphorylated to act as inhibitors^110–112^. Consequently, some high-confidence interactions in our atlas may represent inhibitory or scaffolding relationships rather than enzymatic substrate modifications. Finally, because the atlas is based on a defined confidence threshold, it may exclude genuine low-affinity interactions, and the direct extrapolation of specific findings to other systems requires further system-specific validation.

Future directions include integration with quantitative phosphoproteomics, transcriptomic expression profiles, and genetic interaction maps to refine functional annotation. Extending AFM + iLIS to ubiquitin ligases, phosphatases, and additional protein families will test generality and accelerate discovery of non-canonical regulatory interfaces. The newly identified docking sites—particularly within IDRs—offer untapped opportunities for allosteric and degradation-tag inhibitor design, complementing classical active site-targeting strategies.

By charting both classical and unconventional kinase docking landscapes at unprecedented scale, our atlas empowers hypothesis-driven exploration of signaling regulation and provides a versatile platform for the rational development of next-generation kinase modulators. Ultimately, by providing a structural blueprint of the kinome, this work demonstrates how predictive structural biology is evolving from a tool for understanding individual proteins into an approach for the rational modulation of entire cellular networks.

## Methods

### AlphaFold-Multimer prediction

For AFM predictions, we employed LocalColabFold v.1.5.5^113^, which integrates AlphaFold-Multimer (AFM) v.2.3.1. MSAs were generated using colabfold_search against the UniRef30 (version 2202) and mmseqs2/14-7e284 structural databases. All computations were performed on the Harvard O2 high-performance computing cluster and the Longwood high-performance compute cluster using a variety of NVIDIA GPUs (TeslaV100, RTX6000, RTX8000, A40, A100, L40S, and H100). Each prediction generated five models with five recycling. Interaction confidence was assessed using the integrated Local Interaction Score (iLIS), with a high-confidence threshold of iLIS ≥ 0.223 established at a 10% FDR based on a comprehensive benchmark against curated positive and control reference sets (see **Supplementary Text 1-4**). iLIS analysis code and benchmarking thresholds based on literature-based datasets are available at https://github.com/flyark/AFM-LIS.

### Interaction fingerprint clustering

To group a target protein’s interaction partners based on their binding patterns, we developed a pipeline to generate and cluster “interaction fingerprints.” For each target protein, a set of high-confidence binding partners (iLIS ≥ 0.223) was used as input.

First, a binary contact matrix was constructed where each row represented an interacting partner and each column represented a residue position on the target protein. A value of 1 was assigned if a partner made contact with a specific target protein residue (as defined by the contact-filtered Local Interaction Residues, or cLIRs), and 0 otherwise.

These binary fingerprints were then subjected to agglomerative hierarchical clustering using the SciPy library^114^. The similarity between fingerprints was measured using the cosine distance, and clusters were merged using the average linkage (UPGMA) algorithm. To determine the optimal number of clusters for each target protein automatically, we tested a range of cluster numbers (from 2 to 30) and calculated the Silhouette Score for each partition using the scikit-learn library^115^. This score measures the quality of the clustering, and the number of clusters that maximized this score was chosen as the optimal grouping. The resulting clusters, representing distinct binding modes, were visualized using a clustermap (seaborn.clustermap)^116^ where partners are ordered by the clustering dendrogram.

### Binding-defined kinase tree generation

The binding-defined kinase tree was generated in R from a kinase co-occurrence matrix derived from a stability analysis of 10,800 clustering runs (see **Supplementary Text 7**). The similarity matrix was converted to a distance matrix and subjected to agglomerative hierarchical clustering (complete linkage). The final clusters were defined using the Dynamic Tree Cutting algorithm (cutreeDynamic R package)^117^. The resulting dendrogram was visualized using the ggtree package^118^.

### Sequence-based kinase tree generation

To generate a phylogenetic tree based on sequence homology (as shown in Figure 4D), kinase domain sequences from *Drosophila melanogaster* were obtained, with family classifications (e.g., AGC, CMGC) based on annotations from KinBase^55^ and FlyBase^119^. The analysis was performed in R. First, a multiple sequence alignment of the kinase domain amino acid sequences was generated using the Muscle algorithm^120^, implemented in the msa package^121^. The resulting alignment was then used to calculate a maximum likelihood distance matrix using the JTT (Jones-Taylor-Thornton) model of amino acid substitution ^122^ via the phangorn package^123^. From this distance matrix, a phylogenetic tree was constructed using the Neighbor-Joining algorithm^124^, as implemented in the ape package^125^. Finally, all kinases belonging to the “Atypical” class were pruned from the tree to focus the visualization on the major kinase families. The final phylogeny was plotted as an unrooted tree using the ggtree package^118^.

### Validation of predicted protein-protein interactions using external datasets

To validate the predicted human kinase–transcription factor interactions, the atlas was cross-referenced against two independent, large-scale datasets. For physical and genetic Interactions, known protein-protein and genetic interactions were sourced from the BioGRID database (Version 4.4.245)^47^. The datasets for *Homo sapiens* and *Drosophila melanogaster* were downloaded and integrated for comparison with our predictions. To identify functional co-dependency, functional relationships were assessed using data from the Cancer Dependency Map (DepMap, Public 25Q2)^48^. High-confidence structural predictions from our atlas were compared against the top 100 functionally co-dependent gene partners (ranked by absolute correlation) for each corresponding kinase or transcription factor in the DepMap CRISPR screen dataset.

### Comparison with experimentally derived PWMs

Computationally derived Position Weight Matrices (PWMs) were compared to experimentally determined motifs from Johnson et al., 2023^4^. The normalized and scaled PSSM data (ser_thr_all_norm_scaled_matrice), raw matrix data (ser_thr_all_raw_matrices), and kinase metadata were extracted from the supplementary materials of the publication for direct comparison and visualization. A mapping between the gene names used in our study and the “Matrix_name” identifiers was created to ensure accurate correspondence between predicted and experimental motifs. Sequence logos for human PWMs were generated using logomaker pagage in R^126^.

### Data analysis and visualization

Statistical analyses, including Receiver Operating Characteristic (ROC) curve analysis, were performed using the scikit-learn^115^ and SciPy^114^ libraries in Python. Figures were generated using Matplotlib^127^, Seaborn^116^, and logomaker^128^ in Python, or the ggtree package^118^ in R. Protein structures and electrostatic potentials were visualized in UCSF ChimeraX v1.9^129^. Electrostatic surfaces were calculated in ChimeraX using the coulomb command with a rdbu-7 palette and a charge range of -12 to 12.

### Cell culture and transfection

*Drosophila* S2R+ cells (DGRC, Cat# 150) were maintained at 25°C in Schneider’s Drosophila Medium (Thermo Fisher Scientific, Cat# 21720-024) supplemented with 10% Fetal Bovine Serum (Sigma, Cat# A3912) and 50 U/ml Penicillin-Streptomycin (Thermo Fisher Scientific, Cat# 15070-063). For all experiments, cells were transfected in 6-well plates using the Effectene Transfection Reagent (Qiagen, Cat# 301427) according to the manufacturer’s protocol.

### MBP fusion protein expression and purification

Expression vectors were constructed using synthetic DNA fragments encoding Ftz and Gsb homeodomain sequences with and without mutations (Integrated DNA Technologies) cloned into the pMal-c6T plasmid (New England Biolabs, Cat# N0378S). BL21(DE3) competent E. coli (New England Biolabs, Cat# C2527I) were transformed and cultured overnight. Cultures were diluted 1:100 into LB medium supplemented with 100 μg/mL ampicillin and 0.1% glucose, grown at 37°C to an OD600 of 0.6, and protein expression was induced with 0.3 mM IPTG overnight at 15°C. Bacterial pellets were collected by centrifugation, resuspended in PBS supplemented with 1 mM DTT and 2× Protease Inhibitor Cocktail (Sigma-Aldrich, Cat# S8830-20TAB), and lysed by sonication using a BioRuptor (Diagenode) with 20 cycles of 30 sec ON/30 sec OFF at 4°C. Lysates were supplemented with 1% Triton X-100 (final concentration) and incubated for 30 min at 4°C. Cleared lysates were obtained by centrifugation at 20,000 × g for 10 min at 4°C and incubated with amylose resin (New England Biolabs, Cat# E8021L) for 2 hours at 4°C. To remove DNA bound to the homeodomain fragments, the resins were washed three times with PBS containing 1 mM DTT and 1 M NaCl, followed by two washes with PBS containing 1 mM DTT. MBP-tagged proteins were eluted in 10 mM maltose in PBS and stored at −80°C after dialysis in PBS containing 1 mM DTT.

### MBP pulldown assays

For pulldown experiments, 15 μL of Amylose Magnetic Beads (New England Biolabs, Cat# E8035S) were incubated with purified MBP fusion proteins (5 μg) for 1 hour at 4°C with rotation, then washed three times with CHAPS lysis buffer (Boston BioProducts, Cat# BP-114) supplemented with 1 M NaCl. Beads were then incubated overnight at 4°C with gentle rotation in S2R+ cell lysates expressing aPKC-VHH05-3xHA or Sqa-3xHA prepared in CHAPS lysis buffer supplemented with 2× protease and phosphatase inhibitor cocktails (Thermo Fisher Scientific, Cat# A32961). Following incubation, beads were washed six times with CHAPS lysis buffer, and bound proteins were eluted by boiling at 95°C for 5 min in SDS sample buffer. Eluates were separated by SDS-PAGE using stain-free gels (Bio-Rad, Cat# 4568096) and analyzed by western blotting. Immobilized MBP proteins were detected using stain-free gel imaging. Co-purified kinases were detected using NbVHH05-hIgG nanobody (Addgene, Cat# 171565)(1:100 dilution; in-house prepared)^130^ or rat anti-HA 3F10 antibody (1:1000 dilution; Millipore Sigma, Cat# 11867423001) with corresponding HRP-conjugated secondary antibodies (anti-Human-HRP, BioLegend, Cat# 410603; anti-Rat-HRP, Jackson ImmunoResearch, Cat# 112-035-062).

### *In vitro* phosphorylation assays

MBP fusion proteins (5 μg) were immobilized on Amylose Magnetic Beads as described above and incubated with S2R+ cell lysates expressing aPKC-VHH05-3xHA or empty vector control for 3 hours at 4°C with gentle rotation. Beads were washed six times with CHAPS lysis buffer, and proteins were eluted by boiling at 95°C for 5 min. Phosphorylation was detected by western blot using rabbit anti-phospho-Ser/Thr antibody (1:1000 dilution; Cell Signaling Technology, Cat# 9631S) with HRP-conjugated anti-rabbit secondary antibody (Jackson ImmunoResearch, Cat# 111-035-003). MBP loading was confirmed using stain-free gel imaging (Bio-Rad ChemiDoc MP imaging system).

### Cloning and expression

Expression vectors were constructed using cDNA from the FlyBi ORF collection ^35^ or synthetic DNA fragments (IDT or Twist Bioscience). For validation of allosteric hotspots, transcription factor fragments identified in the AFM screens against Pdk1 and Dsor1 were synthesized and cloned into the pMT vector with N-terminus 127D01 tag and C-terminus BFP tag (pMT-127D01-Binder-BFP).

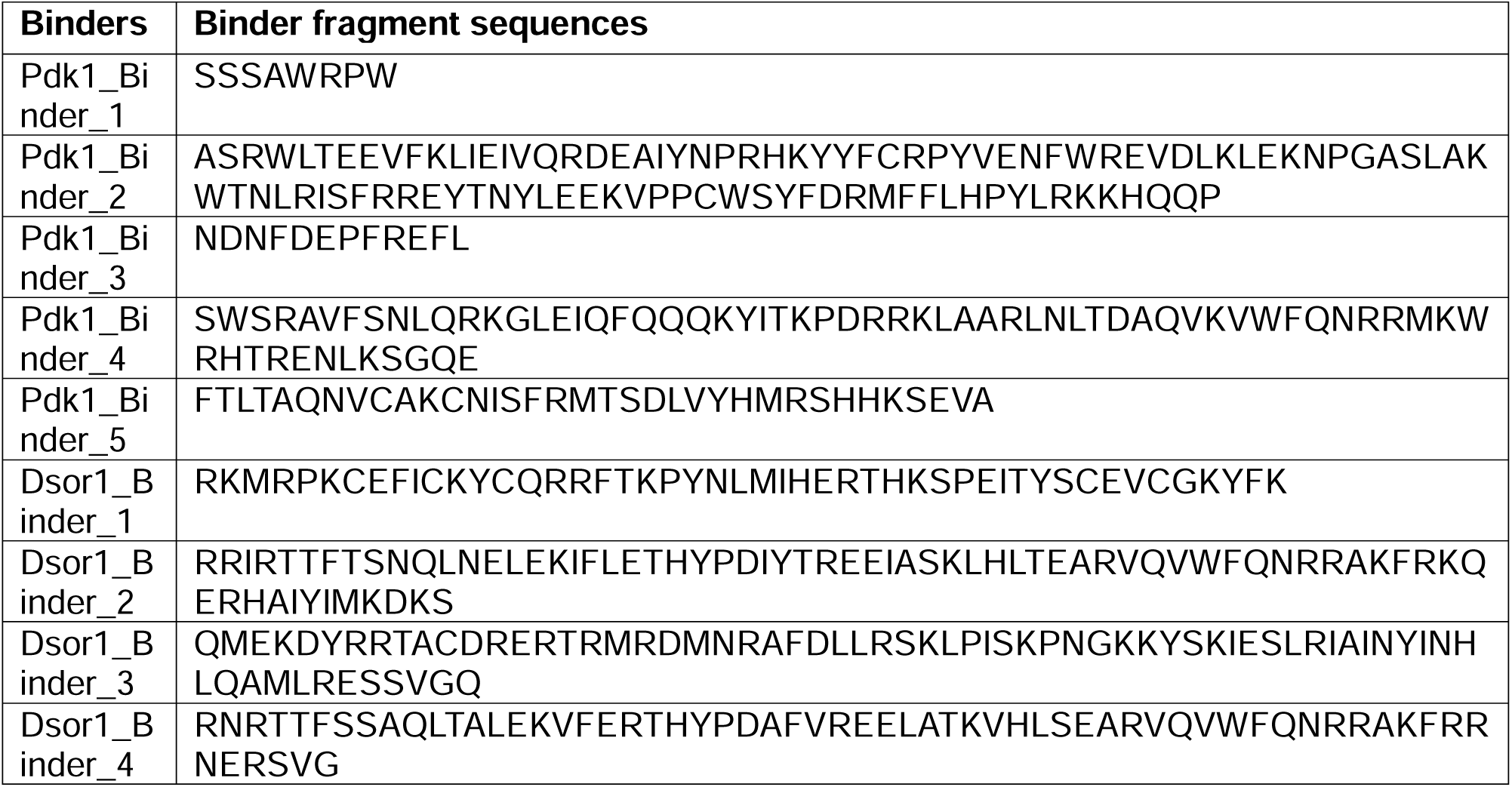

For the Hnf4 analysis, DNA fragments for full-length Hnf4 and the Hnf4[178-493] fragment were synthesized and cloned into a pMT vector for N-terminal GFP tagging (pMT-CMP-v1-GFP-Hnf4-full and pMT-CMP-v1-GFP-Hnf4-[178-493]). The Sgg FlyBi clone was cloned into the pAWH vector (DGRC, Cat# 1096) for C-terminal 3xHA tagging (pAWH-sgg).

Expression from vectors containing the pMT promoter was induced with 500 µM copper sulfate (CuSO4). Other constructs were expressed constitutively from pAct promoter.

### Co-immunoprecipitation and western blotting

For co-IP experiments, transfected cells were harvested, washed twice in PBS, and lysed on ice in CHAPS lysis buffer (Boston BioProducts, Cat# BP-114) supplemented with 2x protease and phosphatase inhibitor cocktails (Thermo Fisher Scientific, Cat# A32961). Cell lysates were cleared by centrifugation at 20,000 x g for 10 minutes at 4°C.

Cleared lysates were incubated with nanobodies coupled to magnetic beads overnight at 4°C with rotation. Proteins tagged with GFP were immunoprecipitated using an in-house purified anti-GFP nanobody (NbGFP-ALFA-His) conjugated to ALFA-tag magnetic resin (NanoTag Biotechnologies, Cat# N1512). Proteins tagged with mCherry were immunoprecipitated using mCherry-Trap magnetic resin (ChromoTek, Cat# rtma). Following incubation and washes, bound proteins were eluted, separated by SDS-PAGE (Bio-Rad, Cat# 4568096), and analyzed by western blot.

Primary antibodies used were: rabbit anti-GFP (Invitrogen, Cat# A-6455), rat anti-HA (Roche, Cat# 11867423001), rabbit anti-P-Akt (T342) (PhosphoSolutions, Cat# p104-342), rabbit anti-Akt (Cell Signaling Technology, Cat# 9272S), rabbit anti-P-ERK1/2 (Cell Signaling Technology, Cat# 4370S), and rabbit anti-ERK1/2 (Cell Signaling Technology, Cat# 4695S). An anti-Actin-Rhodamine antibody (Bio-Rad, Cat# 12004163) was used as a loading control. HRP-conjugated anti-rabbit (Jackson ImmunoResearch, Cat# 111-035-003) and anti-rat secondary antibodies (Jackson ImmunoResearch, Cat# 112-035-062) were used for detection with Pico Plus (Thermo Scientific, Cat# 34580) or Femto ECL chemiluminescent substrates (Thermo Scientific, Cat# 34095). Images were acquired using a Bio-Rad ChemiDoc MP imaging system.

### Lithium chloride treatment

To assess protein stability, S2R+ cells were transfected with a pMT-CMP-v1-GFP-Hnf4-full plasmid. Following a 24-hour induction with 500 µM CuSO4, the GSK3 inhibitor lithium chloride (LiCl) was added to the media for an additional 16 hours. Cells were then collected and processed for western blot analysis.

### Phosphoproteomics

S2R+ cells were transfected with a Hnf4[178-493] expression vector (pMT-CMP-v1-GFP-Hnf4-[178-493]). After induction with 500 µM CuSO4, cells were treated with either a control buffer or 10 mM LiCl for 16 hours. The GFP-Hnf4[178-493] fragment was immunoprecipitated from cleared cell lysates using anti-GFP nanobody conjugated to ALFA-tag magnetic resin. Eluates by ALFA peptide were separated by SDS-PAGE, and the gel bands corresponding to the molecular weight of the Hnf4 fragment were excised and submitted for phosphorylation site mapping in the Mass Spectrometry Facility at Beth Israel Deaconess Medical Center.

### *Drosophila* stocks and husbandry

All experiments were conducted using female flies due to their larger oenocyte size, which facilitated dissection. For temperature-sensitive experiments, fly crosses were set up and maintained under a 12:12 hour light:dark cycle at 18°C. Crosses were discarded after 5 days to maintain consistent population density. Following eclosion, adult flies were collected and kept at 18°C for 2–3 days. *w; Hnf4-127D01; PromE-GAL4, Tub-GAL80^TS^* flies were crossed with following Bloomington Drosophila Stock Center RNAi lines: *par-1* (#32410), *gish* (#36719), *sgg* (#38293), *Lk6* (#60003), *Pdk1* (#34936), *Cdk12* (#34838), *nmo* (#60016), *Tao* (#34881), *Drak* (#44102), *Abl* (#61170), *Dyrk2* (#57294), *Pkc98E* (#44074), *Src42A* (#44039), *PAN3* (#57037), *dop* (#76064), *Pkn* (#55876), *CG17528* (#42575), and *Adck5* (#57406).

### *In vivo* immunostaining

For nanobody immunostaining, adult fly abdomens were dissected in PBS at room temperature, the fat body was removed, and tissues were fixed in 4% paraformaldehyde in PBS for 30 min. Samples were washed 3 × 5 min in PBST (PBS + 0.1% Triton X-100) and blocked in PBST + 5% normal goat serum (NGS) for 1 h. The anti-ALFA nanobody Nb127D01-ALFA (0.2 mg/mL stock) was diluted 1:500 in PBST + 5% NGS and applied for 1.5 h at room temperature with gentle agitation, followed by 3 × 5 min washes in PBST. Secondary antibody Alexa Fluor 594 anti-Alpaca (Jackson ImmunoResearch, Cat# 128-585-232) was diluted 1:400 in PBST + 5% NGS and incubated with samples for 1 h at room temperature. After PBST washes and DAPI counterstaining, tissues were mounted in VECTASHIELD Antifade Mounding Medium (Vector Laboratories, Cat# H-1000-10) and imaged on an Olympus VS200 slide scanner or a Zeiss Axio Observer Z1 confocal microscope.

### Oil Red O staining

Oil Red O staining was performed as previously described<u>(Gutierrez *et al.*, 2007)</u> with minor modifications. Adult flies were dissected in ice-cold PBS, and fat bodies were removed by gentle aspiration (“liposuction”). Tissues were fixed in 4% paraformaldehyde for 20 min at room temperature, rinsed twice in distilled water, and stained with freshly prepared Oil Red O. For the stain, Oil Red O (Sigma-Aldrich, Cat# O0625) was dissolved in isopropanol to 0.1% (w/v) to prepare a stock solution; the working solution was made immediately before use by mixing 3 mL of stock with 2 mL of distilled water and filtering through a 0.2-µm syringe filter. Samples were incubated in the working solution for 25 min, briefly rinsed in PBS, and mounted in VECTASHIELD medium for imaging. Images were acquired on an Olympus VS200 slide scanner equipped with a color camera (2448 × 2048 pixels; 3.45-µm pixel size).

## Supporting information

SF 8_cluster 1

## Acknowledgements

We are grateful to the Research Computing Group at Harvard Medical School for GPU access in the Harvard O2 high-performance computing cluster and the Longwood high-performance compute cluster and the Microscopy Resources on North Quad (MicRoN) core at Harvard Medical School for confocal microscopy access. We thank Mr. Jonathan Rodiger, Mr. Aram Comjean and Mr. Austin Veal for help to develop the FlyPredictome portal and Dr. John M. Asara at the Beth Israel Deaconess Medical Center for his help performing mass spectrometry.

This research was supported by the National Institutes of Health through grants P01 CA120964 (to N.P. and L.C.C.), P41 GM132087 and R01 AR057352 (to N.P.), and R35 CA197588 and P01 CA117969 (to L.C.C.). Further funding was provided by a Claudia Adams Barr Program for Cancer Research award (to J.L.J.) and a Postdoctoral Fellowship from the National Research Foundation of Korea (NRF) funded by the Ministry of Education (2021R1A6A3A14039622 to A.K.). N.P. is an investigator of the Howard Hughes Medical Institute.

During the preparation of this manuscript, the authors used OpenAI’s ChatGPT and Google’s Gemini to assist in generating custom Python/R scripts for the analysis of prediction results and to improve the readability and language of the text. After using this tool, the authors reviewed and edited the content as needed and take full responsibility for the publication.

This article is subject to HHMI’s Open Access to Publications policy. HHMI lab heads have previously granted a nonexclusive CC BY 4.0 license to the public and a sublicensable license to HHMI in their research articles. Pursuant to those licenses, the author-accepted manuscript of this article can be made freely available under a CC BY 4.0 license immediately upon publication.

## Author Contributions

A.K. and N.P. conceived this study. A.K. developed the bioinformatic methodology and wrote the original manuscript. The experimental investigation was conducted as follows: the *in vivo* RNAi screen was performed by K.H. and K.W.; the kinase profiling for human GRK7 was performed by J.L.J., T.M.Y., and L.C.C.; and all remaining experiments were performed by A.K. The final manuscript was reviewed and edited by A.K., Y.H., and N.P., with input from all authors.

## Conflict of Interest

L.C.C. is a founder and member of the board of directors of Agios Pharmaceuticals and is a founder and receives research support from Petra Pharmaceuticals; is listed as an inventor on a patent (WO2019232403A1, Weill Cornell Medicine) for combination therapy for PI3K-associated disease or disorder, and the identification of therapeutic interventions to improve response to PI3K inhibitors for cancer treatment; is a co-founder and shareholder in Faeth Therapeutics; has equity in and consults for Cell Signaling Technologies, Volastra, Larkspur and 1 Base Pharmaceuticals; and consults for Loxo-Lilly. T.M.Y. is a co-founder of DeStroke. J.L.J. has received consulting fees from Scorpion Therapeutics and Volastra Therapeutics. The remaining authors declare no competing interests.

## Data Availability Statement

The complete fly kinase–TF atlas, including iLIS scores and models, is available for interactive exploration at the FlyPredictome portal (https://www.flyrnai.org/tools/fly_predictome). All supplementary datasets generated in this study, including the residue-level interaction data (**Supplementary Data 1** and **3**), the complete PWMs for the human and fly screens (**Supplementary Data 2** and **4**), the comprehensive catalog of interaction hotspots (**Supplementary Data 5**), and the multi-page **Supplementary Figure 8** (containing sequence logos for all fly kinase clusters) have been deposited in the Zenodo repository and are publicly available under DOI: 10.5281/zenodo.17314335.

## Notes

https://www.flyrnai.org/tools/fly_predictome/web/

https://zenodo.org/records/17314335

### Summary of Updates

Figures with high DPI are updated.

## References

1. Songyang, Z. et al. Use of an oriented peptide library to determine the optimal substrates of protein kinases. Curr. Biol. CB 4, 973–982 (1994).

2. Nishikawa, K., Toker, A., Johannes, F. J., Songyang, Z. & Cantley, L. C. Determination of the specific substrate sequence motifs of protein kinase C isozymes. J. Biol. Chem. 272, 952–960 (1997).

3. Brinkworth, R. I., Breinl, R. A. & Kobe, B. Structural basis and prediction of substrate specificity in protein serine/threonine kinases. Proc. Natl. Acad. Sci. 100, 74–79 (2003).

4. Johnson, J. L. et al. An atlas of substrate specificities for the human serine/threonine kinome. Nature 613, 759–766 (2023).

5. Yaron-Barir, T. M. et al. The intrinsic substrate specificity of the human tyrosine kinome. Nature 629, 1174–1181 (2024).

6. Reményi, A., Good, M. C. & Lim, W. A. Docking interactions in protein kinase and phosphatase networks. Curr. Opin. Struct. Biol. 16, 676–685 (2006).

7. Goldsmith, E. J., Akella, R., Min, X., Zhou, T. & Humphreys, J. M. Substrate and Docking Interactions in Ser/Thr Protein Kinases. Chem. Rev. 107, 5065–5081 (2007).

8. Miller, C. J. & Turk, B. E. Homing in: Mechanisms of Substrate Targeting by Protein Kinases. Trends Biochem. Sci. 43, 380–394 (2018).

9. Jumper, J. et al. Highly accurate protein structure prediction with AlphaFold. Nature 596, 583–589 (2021).

10. Evans, R. et al. Protein complex prediction with AlphaFold-Multimer. 2021.10.04.463034 Preprint at 10.1101/2021.10.04.463034 (2022).

11. Abramson, J. et al. Accurate structure prediction of biomolecular interactions with AlphaFold 3. Nature 630, 493–500 (2024).

12. Baek, M. et al. Accurate prediction of protein structures and interactions using a three-track neural network. Science 373, 871–876 (2021).

13. Baek, M. et al. Efficient and accurate prediction of protein structure using RoseTTAFold2. 2023.05.24.542179 Preprint at 10.1101/2023.05.24.542179 (2023).

14. Humphreys, I. R. et al. Computed structures of core eukaryotic protein complexes. Science 374, eabm4805 (2021).

15. Durham, J., Zhang, J., Humphreys, I. R., Pei, J. & Cong, Q. Recent advances in predicting and modeling protein–protein interactions. Trends Biochem. Sci. 48, 527–538 (2023).

16. Burke, D. F. et al. Towards a structurally resolved human protein interaction network. Nat. Struct. Mol. Biol. 30, 216–225 (2023).

17. Humphreys, I. R. et al. Protein interactions in human pathogens revealed through deep learning. Nat. Microbiol. 9, 2642–2652 (2024).

18. Yoon, M. S., Bae, B., Kim, K., Park, H. & Baek, M. Deep learning methods for proteome-scale interaction prediction. Curr. Opin. Struct. Biol. 90, 102981 (2025).

19. Schmid, E. W. & Walter, J. C. Predictomes, a classifier-curated database of AlphaFold-modeled protein-protein interactions. Mol. Cell 85, 1216–1232.e5 (2025).

20. Laman Trip, D. S., et al. A tissue-specific atlas of protein–protein associations enables prioritization of candidate disease genes. Nat. Biotechnol. 1–14 (2025) doi:10.1038/s41587-025-02659-z.

21. Yuan, R., Zhang, J., Zhou, J. & Cong, Q. Recent progress and future challenges in structure-based protein-protein interaction prediction. Mol. Ther. 33, 2252–2268 (2025).

22. Zhang, J. et al. Predicting protein-protein interactions in the human proteome. Science 0, eadt1630 (2025).

23. Schmid, E. W. et al. Proteome-wide in silico screening for human protein-protein interactions. 2025.11.10.687652 Preprint at 10.1101/2025.11.10.687652 (2025).

24. Wright, P. E. & Dyson, H. J. Intrinsically Disordered Proteins in Cellular Signaling and Regulation. Nat. Rev. Mol. Cell Biol. 16, 18–29 (2015).

25. Holehouse, A. S. & Kragelund, B. B. The molecular basis for cellular function of intrinsically disordered protein regions. Nat. Rev. Mol. Cell Biol. 25, 187–211 (2024).

26. Ruff, K. M. et al. Molecular grammars of predicted intrinsically disordered regions that span the human proteome. Cell S0092-8674(25)01191–2 (2025) doi:10.1016/j.cell.2025.10.019.

27. Whitmarsh, A. J. Regulation of gene transcription by mitogen-activated protein kinase signaling pathways. Biochim. Biophys. Acta BBA - Mol. Cell Res. 1773, 1285–1298 (2007).

28. Weidemüller, P., Kholmatov, M., Petsalaki, E. & Zaugg, J. B. Transcription factors: Bridge between cell signaling and gene regulation. PROTEOMICS 21, 2000034 (2021).

29. Tesei, G. et al. Conformational ensembles of the human intrinsically disordered proteome. Nature 626, 897–904 (2024).

30. Datta, R. R., Akdogan, D., Tezcan, E. B. & Onal, P. Versatile roles of disordered transcription factor effector domains in transcriptional regulation. FEBS J. 292, 3014–3033 (2025).

31. Bryant, P., Pozzati, G. & Elofsson, A. Improved prediction of protein-protein interactions using AlphaFold2. Nat. Commun. 13, 1265 (2022).

32. Lee, C. Y. et al. Systematic discovery of protein interaction interfaces using AlphaFold and experimental validation. Mol. Syst. Biol. 20, 75–97 (2024).

33. Kim, A.-R. et al. Enhanced Protein-Protein Interaction Discovery via AlphaFold-Multimer. 2024.02.19.580970 Preprint at 10.1101/2024.02.19.580970 (2024).

34. Braun, P. et al. An experimentally derived confidence score for binary protein-protein interactions. Nat. Methods 6, 91–97 (2009).

35. Tang, H.-W. et al. Next-generation large-scale binary protein interaction network for Drosophila melanogaster. Nat. Commun. 14, 2162 (2023).

36. Yu, H. et al. High-quality binary protein interaction map of the yeast interactome network. Science 322, 104–110 (2008).

37. Kumar, M. et al. ELM—the Eukaryotic Linear Motif resource—2024 update. Nucleic Acids Res. 52, D442–D455 (2024).

38. Varga, J. K., Ovchinnikov, S. & Schueler-Furman, O. actifpTM: a refined confidence metric of AlphaFold2 predictions involving flexible regions. Bioinforma. Oxf. Engl. 41, (2025).

39. Dunbrack, R. L. J. Rēs ipSAE loquunt: What’s wrong with AlphaFold’s ipTM score and how to fix it. Preprint at 10.1101/2025.02.10.637595 (2025).

40. Harkiolaki, M. et al. Structural basis for SH3 domain-mediated high-affinity binding between Mona/Gads and SLP-76. EMBO J. 22, 2571–2582 (2003).

41. Kussie, P. H. et al. Structure of the MDM2 oncoprotein bound to the p53 tumor suppressor transactivation domain. Science 274, 948–953 (1996).

42. An, W. G. et al. Stabilization of wild-type p53 by hypoxia-inducible factor 1α. Nature 392, 405–408 (1998).

43. Ravi, R. et al. Regulation of tumor angiogenesis by p53-induced degradation of hypoxia-inducible factor 1alpha. Genes Dev. 14, 34–44 (2000).

44. Morcillo, P., Rosen, C., Baylies, M. K. & Dorsett, D. Chip, a widely expressed chromosomal protein required for segmentation and activity of a remote wing margin enhancer in Drosophila. Genes Dev. 11, 2729–2740 (1997).

45. Torigoi, E. et al. Chip interacts with diverse homeodomain proteins and potentiates Bicoid activity in vivo. Proc. Natl. Acad. Sci. 97, 2686–2691 (2000).

46. van Meyel, D. J. et al. Chip and Apterous Physically Interact to Form a Functional Complex during *Drosophila* Development. Mol. Cell 4, 259–265 (1999).

47. Oughtred, R. et al. The BioGRID database: A comprehensive biomedical resource of curated protein, genetic, and chemical interactions. Protein Sci. 30, 187–200 (2021).

48. Arafeh, R., Shibue, T., Dempster, J. M., Hahn, W. C. & Vazquez, F. The present and future of the Cancer Dependency Map. Nat. Rev. Cancer 25, 59–73 (2025).

49. Liu, S., Sun, J.-P., Zhou, B. & Zhang, Z.-Y. Structural basis of docking interactions between ERK2 and MAP kinase phosphatase 3. Proc. Natl. Acad. Sci. 103, 5326–5331 (2006).

50. Peti, W. & Page, R. Molecular basis of MAP kinase regulation. Protein Sci. Publ. Protein Soc. 22, 1698–1710 (2013).

51. Francis, D. M., Koveal, D., Tortajada, A., Page, R. & Peti, W. Interaction of Kinase-Interaction-Motif Protein Tyrosine Phosphatases with the Mitogen-Activated Protein Kinase ERK2. PLoS ONE 9, e91934 (2014).

52. McInnes, L., Healy, J. & Melville, J. UMAP: Uniform Manifold Approximation and Projection for Dimension Reduction. Preprint at 10.48550/arXiv.1802.03426 (2020).

53. Hanks, S. K., Quinn, A. M. & Hunter, T. The protein kinase family: conserved features and deduced phylogeny of the catalytic domains. Science 241, 42–52 (1988).

54. Manning, G., Whyte, D. B., Martinez, R., Hunter, T. & Sudarsanam, S. The protein kinase complement of the human genome. Science 298, 1912–1934 (2002).

55. Manning, G., Plowman, G. D., Hunter, T. & Sudarsanam, S. Evolution of protein kinase signaling from yeast to man. Trends Biochem. Sci. 27, 514–520 (2002).

56. Asai, D. et al. Peptide substrates for G protein-coupled receptor kinase 2. FEBS Lett. 588, 2129–2132 (2014).

57. Maier, D., Cheng, S., Faubert, D. & Hipfner, D. R. A Broadly Conserved G-Protein-Coupled Receptor Kinase Phosphorylation Mechanism Controls Drosophila Smoothened Activity. PLOS Genet. 10, e1004399 (2014).

58. Eyers, P. A., Keeshan, K. & Kannan, N. Tribbles in the 21st Century: The Evolving Roles of Tribbles Pseudokinases in Biology and Disease. Trends Cell Biol. 27, 284–298 (2017).

59. Richmond, L. & Keeshan, K. Pseudokinases: a tribble-edged sword. FEBS J. 287, 4170–4182 (2020).

60. Dobens, L. L., Nauman, C., Fischer, Z. & Yao, X. Control of Cell Growth and Proliferation by the Tribbles Pseudokinase: Lessons from Drosophila. Cancers 13, 883 (2021).

61. Modi, V. & Dunbrack, R. L. A Structurally-Validated Multiple Sequence Alignment of 497 Human Protein Kinase Domains. Sci. Rep. 9, 19790 (2019).

62. Nüsslein-Volhard, C. & Wieschaus, E. Mutations affecting segment number and polarity in Drosophila. Nature 287, 795–801 (1980).

63. Wakimoto, B. T., Turner, F. R. & Kaufman, T. C. Defects in embryogenesis in mutants associated with the antennapedia gene complex of Drosophila melanogaster. Dev. Biol. 102, 147–172 (1984).

64. Carroll, S. B. & Scott, M. P. Localization of the fushi tarazu protein during Drosophila embryogenesis. Cell 43, 47–57 (1985).

65. Hiromi, Y., Kuroiwa, A. & Gehring, W. J. Control elements of the Drosophila segmentation gene fushi tarazu. Cell 43, 603–613 (1985).

66. Krause, H. M. & Gehring, W. J. Stage-specific phosphorylation of the fushi tarazu protein during Drosophila development. EMBO J. 8, 1197–1204 (1989).

67. Dong, J., Hung, L., Strome, R. & Krause, H. M. A phosphorylation site in the Ftz homeodomain is required for activity. EMBO J. 17, 2308–2318 (1998).

68. Qian, Y. Q. et al. The structure of the *Antennapedia* homeodomain determined by NMR spectroscopy in solution: Comparison with prokaryotic repressors. Cell 59, 573–580 (1989).

69. Trelsman, J., Gönczy, P., Vashishtha, M., Harris, E. & Desplan, C. A single amino acid can determine the DNA binding specificity of homeodomain proteins. Cell 59, 553–562 (1989).

70. Banerjee-Basu, S. & Baxevanis, A. D. Molecular evolution of the homeodomain family of transcription factors. Nucleic Acids Res. 29, 3258–3269 (2001).

71. Berger, M. F. et al. Variation in Homeodomain DNA Binding Revealed by High-Resolution Analysis of Sequence Preferences. Cell 133, 1266–1276 (2008).

72. Kock, K. H. et al. DNA binding analysis of rare variants in homeodomains reveals homeodomain specificity-determining residues. Nat. Commun. 15, 3110 (2024).

73. Bruun, J.-A. et al. The third helix of the homeodomain of paired class homeodomain proteins acts as a recognition helix both for DNA and protein interactions. Nucleic Acids Res. 33, 2661–2675 (2005).

74. Zhou, B., Liu, C., Xu, Z. & Zhu, G. Structural basis for homeodomain recognition by the cell-cycle regulator Geminin. Proc. Natl. Acad. Sci. U. S. A. 109, 8931–8936 (2012).

75. Bürglin, T. R. & Affolter, M. Homeodomain proteins: an update. Chromosoma 125, 497–521 (2016).

76. Chi, Y.-I. Homeodomain Revisited: a Lesson from Disease-causing Mutations. Hum. Genet. 116, 433–444 (2005).

77. Faber, E. B. et al. Development of allosteric and selective CDK2 inhibitors for contraception with negative cooperativity to cyclin binding. Nat. Commun. 14, 3213 (2023).

78. Mingione, V. R., Paung, Y., Outhwaite, I. R. & Seeliger, M. A. Allosteric regulation and inhibition of protein kinases. Biochem. Soc. Trans. 51, 373–385 (2023).

79. Wang, B. et al. An overview of kinase downregulators and recent advances in discovery approaches. Signal Transduct. Target. Ther. 6, 423 (2021).

80. Pan, Y. & Mader, M. M. Principles of Kinase Allosteric Inhibition and Pocket Validation. J. Med. Chem. 65, 5288–5299 (2022).

81. Baldi, S. et al. Advancements in Protein Kinase Inhibitors: From Discovery to Clinical Applications. Research 8, 0747 (2025).

82. Biondi, R. M. et al. Identification of a pocket in the PDK1 kinase domain that interacts with PIF and the C-terminal residues of PKA. EMBO J. 19, 979–988 (2000).

83. Biondi, R. M., Kieloch, A., Currie, R. A., Deak, M. & Alessi, D. R. The PIF-binding pocket in PDK1 is essential for activation of S6K and SGK, but not PKB. EMBO J. 20, 4380–4390 (2001).

84. Palanker, L., Tennessen, J. M., Lam, G. & Thummel, C. S. Drosophila HNF4 regulates lipid mobilization and beta-oxidation. Cell Metab. 9, 228–239 (2009).

85. Storelli, G., Nam, H.-J., Simcox, J., Villanueva, C. J. & Thummel, C. S. Drosophila HNF4 directs a switch in lipid metabolism that supports the transition to adulthood. Dev. Cell 48, 200–214.e6 (2019).

86. Almeida-Oliveira, F., Tuthill, B. F., Gondim, K. C., Majerowicz, D. & Musselman, L. P. dHNF4 regulates lipid homeostasis and oogenesis in *Drosophila melanogaster*. Insect Biochem. Mol. Biol. 133, 103569 (2021).

87. Xu, J. et al. Protein visualization and manipulation in Drosophila through the use of epitope tags recognized by nanobodies. eLife 11, e74326 (2022).

88. Huang, K. et al. Lipid metabolism of hepatocyte-like cells supports intestinal tumor growth by promoting tracheogenesis. 2025.04.04.647255 Preprint at 10.1101/2025.04.04.647255 (2025).

89. Han, H.-S., Kang, G., Kim, J. S., Choi, B. H. & Koo, S.-H. Regulation of glucose metabolism from a liver-centric perspective. Exp. Mol. Med. 48, e218–e218 (2016).

90. Khoury, M. et al. Glycogen synthase kinase 3 activity enhances liver inflammation in MASH. JHEP Rep. Innov. Hepatol. 6, 101073 (2024).

91. Fiol, C. J., Mahrenholz, A. M., Wang, Y., Roeske, R. W. & Roach, P. J. Formation of protein kinase recognition sites by covalent modification of the substrate. Molecular mechanism for the synergistic action of casein kinase II and glycogen synthase kinase 3. J. Biol. Chem. 262, 14042–14048 (1987).

92. Frame, S., Cohen, P. & Biondi, R. M. A common phosphate binding site explains the unique substrate specificity of GSK3 and its inactivation by phosphorylation. Mol. Cell 7, 1321–1327 (2001).

93. Beurel, E., Grieco, S. F. & Jope, R. S. Glycogen synthase kinase-3 (GSK3): regulation, actions, and diseases. Pharmacol. Ther. 0, 114–131 (2015).

94. Boyle, W. J. et al. Activation of protein kinase C decreases phosphorylation of c-Jun at sites that negatively regulate its DNA-binding activity. Cell 64, 573–584 (1991).

95. Godemann, R., Biernat, J., Mandelkow, E. & Mandelkow, E.-M. Phosphorylation of tau protein by recombinant GSK-3β: pronounced phosphorylation at select Ser/Thr-Pro motifs but no phosphorylation at Ser262 in the repeat domain. FEBS Lett. 454, 157–164 (1999).

96. Pandey, M. K. & DeGrado, T. R. Glycogen Synthase Kinase-3 (GSK-3)-Targeted Therapy and Imaging. Theranostics 6, 571–593 (2016).

97. Huang, D., Zhou, T., Lafleur, K., Nevado, C. & Caflisch, A. Kinase selectivity potential for inhibitors targeting the ATP binding site: a network analysis. Bioinformatics 26, 198–204 (2010).

98. Zhao, Z. et al. Exploration of Type II Binding Mode: A Privileged Approach for Kinase Inhibitor Focused Drug Discovery? ACS Chem. Biol. 9, 1230–1241 (2014).

99. Attwood, M. M., Fabbro, D., Sokolov, A. V., Knapp, S. & Schiöth, H. B. Trends in kinase drug discovery: targets, indications and inhibitor design. Nat. Rev. Drug Discov. 20, 839–861 (2021).

100. Bennett, N. R. et al. Improving de novo protein binder design with deep learning. Nat. Commun. 14, 2625 (2023).

101. Bryant, P. & Elofsson, A. Peptide binder design with inverse folding and protein structure prediction. Commun. Chem. 6, 229 (2023).

102. Vázquez Torres, S., et al. De novo design of high-affinity binders of bioactive helical peptides. Nature 626, 435–442 (2024).

103. Bhat, S. et al. De novo design of peptide binders to conformationally diverse targets with contrastive language modeling. Sci. Adv. 11, eadr8638 (2025).

104. Rettie, S. A. et al. Accurate de novo design of high-affinity protein-binding macrocycles using deep learning. Nat. Chem. Biol. 1–9 (2025) doi:10.1038/s41589-025-01929-w.

105. Wu, K. et al. Design of intrinsically disordered region binding proteins. Science 389, eadr8063 (2025).

106. Liu, C. et al. Diffusing protein binders to intrinsically disordered proteins. Nature 644, 809–817 (2025).

107. Pacesa, M. et al. One-shot design of functional protein binders with BindCraft. Nature 1–10 (2025) doi:10.1038/s41586-025-09429-6.

108. Muratspahić, E. et al. De novo design of miniprotein agonists and antagonists targeting G protein-coupled receptors. BioRxiv Prepr. Serv. Biol. 2025.03.23.644666 (2025) doi:10.1101/2025.03.23.644666.

109. Li, Q., Helleday, T. & Bryant, P. RareFoldGPCR: Agonist Design Beyond Natural Amino Acids. 2025.10.01.679733 Preprint at 10.1101/2025.10.01.679733 (2025).

110. Taylor, S. S., Buechler, J. A. & Yonemoto, W. cAMP-DEPENDENT PROTEIN KINASE: FRAMEWORK FOR A DIVERSE FAMILY OF REGULATORY ENZYMES. Annu. Rev. Biochem. 59, 971–1005 (1990).

111. Dalton, G. D. & Dewey, W. L. Protein kinase inhibitor peptide (PKI): a family of endogenous neuropeptides that modulate neuronal cAMP-dependent protein kinase function. Neuropeptides 40, 23–34 (2006).

112. Liu, C., Ke, P., Zhang, J., Zhang, X. & Chen, X. Protein Kinase Inhibitor Peptide as a Tool to Specifically Inhibit Protein Kinase A. Front. Physiol. 11, 574030 (2020).

113. Mirdita, M. et al. ColabFold: making protein folding accessible to all. Nat. Methods 19, 679–682 (2022).

114. Virtanen, P. et al. SciPy 1.0: fundamental algorithms for scientific computing in Python. Nat. Methods 17, 261–272 (2020).

115. Pedregosa, F. et al. Scikit-learn: Machine Learning in Python. J Mach Learn Res 12, 2825–2830 (2011).

116. Waskom, M. L. seaborn: statistical data visualization. J. Open Source Softw. 6, 3021 (2021).

117. Langfelder, P., Zhang, B. & Horvath, S. Defining clusters from a hierarchical cluster tree: the Dynamic Tree Cut package for R. Bioinformatics 24, 719–720 (2008).

118. Yu, G., Smith, D. K., Zhu, H., Guan, Y. & Lam, T. T.-Y. ggtree: an r package for visualization and annotation of phylogenetic trees with their covariates and other associated data. Methods Ecol. Evol. 8, 28–36 (2017).

119. Öztürk-Çolak, A. et al. FlyBase: updates to the Drosophila genes and genomes database. Genetics 227, iyad211 (2024).

120. Edgar, R. C. MUSCLE: multiple sequence alignment with high accuracy and high throughput. Nucleic Acids Res. 32, 1792–1797 (2004).

121. Bodenhofer, U., Bonatesta, E., Horejš-Kainrath, C. & Hochreiter, S. msa: an R package for multiple sequence alignment. Bioinforma. Oxf. Engl. 31, 3997–3999 (2015).

122. Jones, D. T., Taylor, W. R. & Thornton, J. M. The rapid generation of mutation data matrices from protein sequences. Bioinformatics 8, 275–282 (1992).

123. Schliep, K. P. phangorn: phylogenetic analysis in R. Bioinformatics 27, 592–593 (2011).

124. Saitou, N. & Nei, M. The neighbor-joining method: a new method for reconstructing phylogenetic trees. Mol. Biol. Evol. 4, 406–425 (1987).

125. Paradis, E. & Schliep, K. ape 5.0: an environment for modern phylogenetics and evolutionary analyses in R. Bioinformatics 35, 526–528 (2019).

126. Wagih, O. ggseqlogo: a versatile R package for drawing sequence logos. Bioinforma. Oxf. Engl. 33, 3645–3647 (2017).

127. Hunter, J. D. Matplotlib: A 2D Graphics Environment. Comput. Sci. Eng. 9, 90–95 (2007).

128. Tareen, A. & Kinney, J. B. Logomaker: beautiful sequence logos in Python. Bioinformatics 36, 2272–2274 (2020).

129. Meng, E. C. et al. UCSF ChimeraX: Tools for structure building and analysis. Protein Sci. 32, e4792 (2023).

130. Kim, A.-R., et al. NanoTag Nanobody Tools for Drosophila In Vitro and In Vivo Studies. Curr. Protoc. 2, e628 (2022).

