## Supplementary figures and images for "A Structure-Guided Kinase–Transcription Factor Interactome Atlas Reveals Docking Landscapes of the Kinome"

### SF 1

#### Supplementary Figure 1

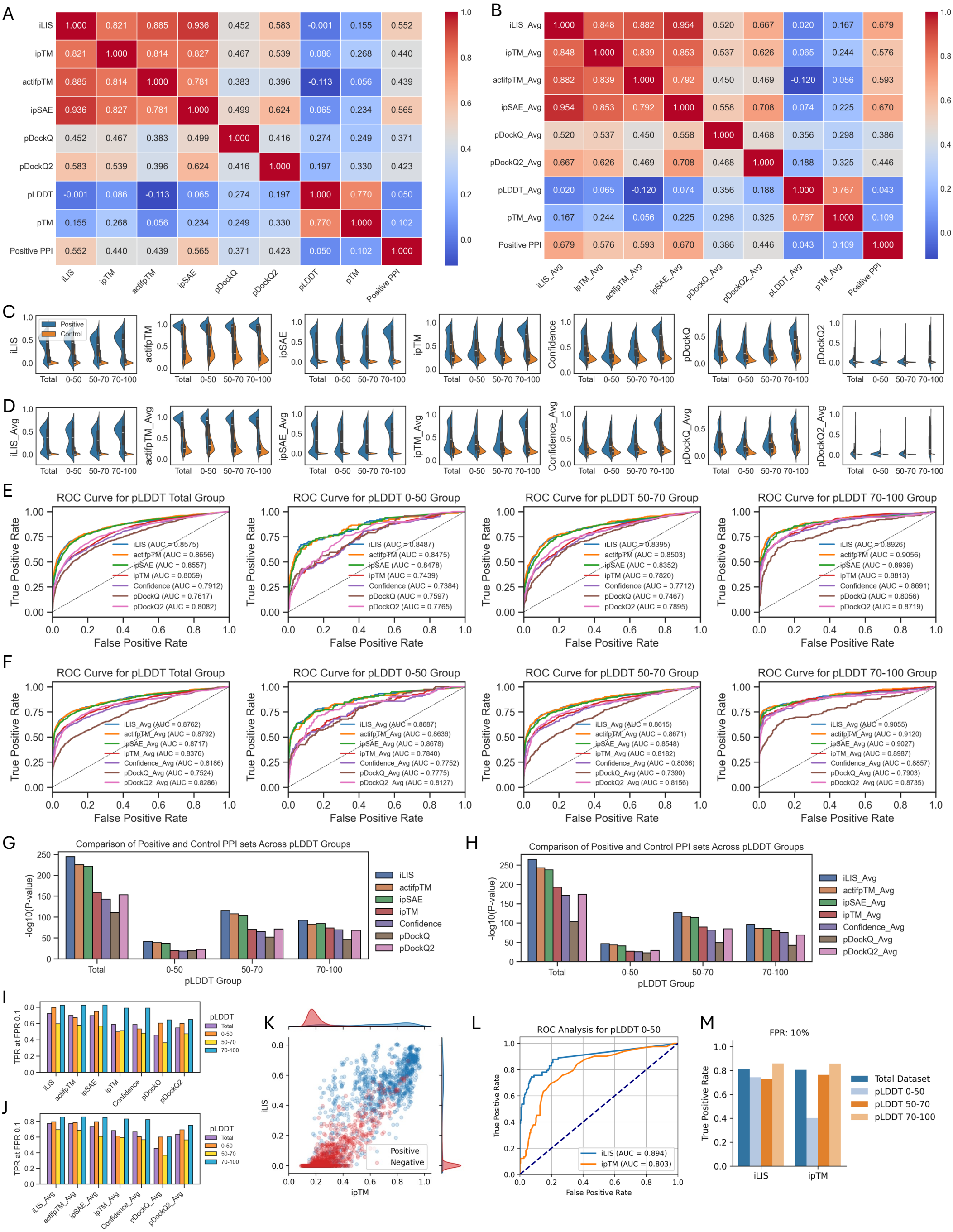

### SF 3

#### Supplementary Figure 3

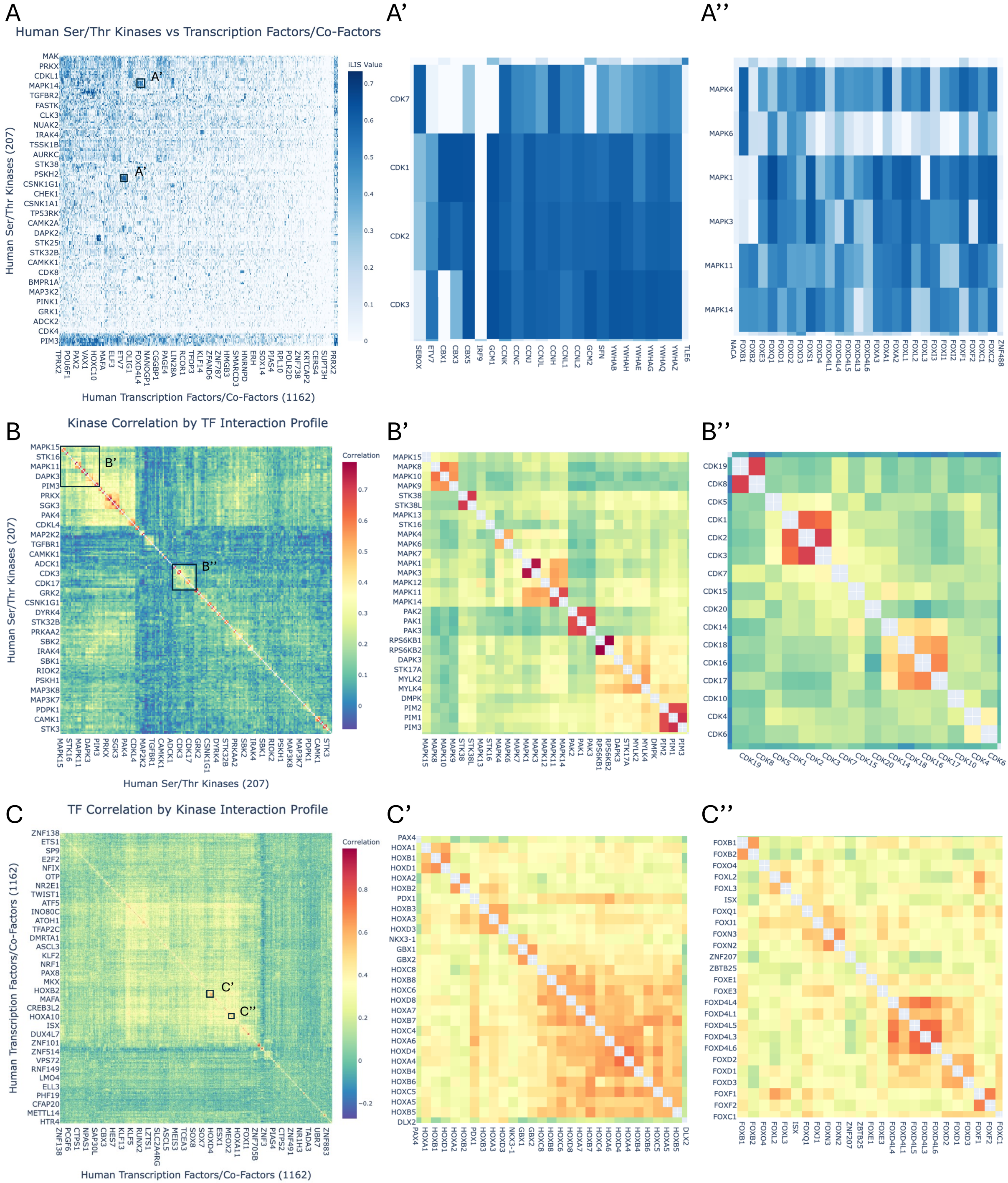

### SF 4A

#### Supplementary Figure 4

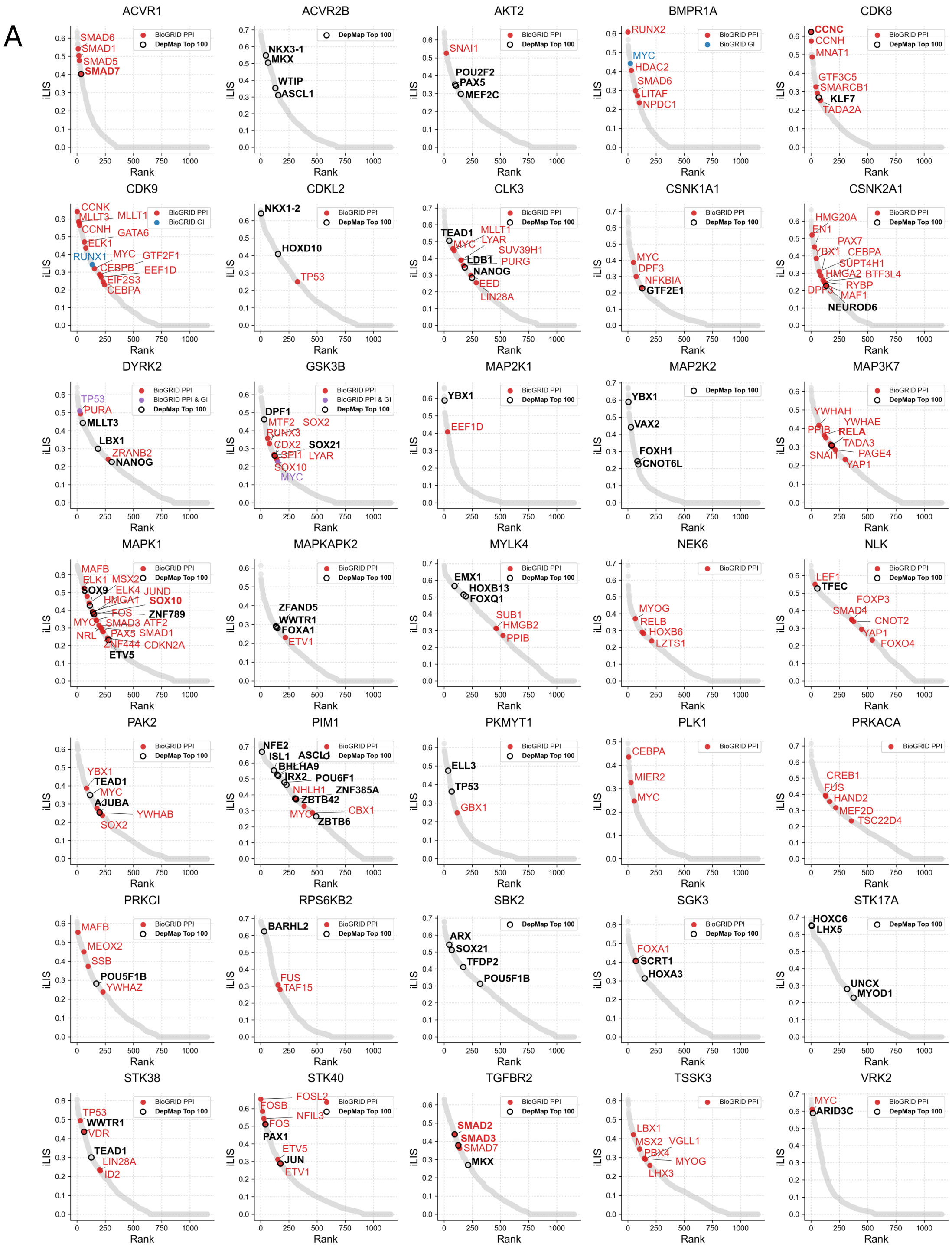

### SF 4B

#### Supplementary Figure 4

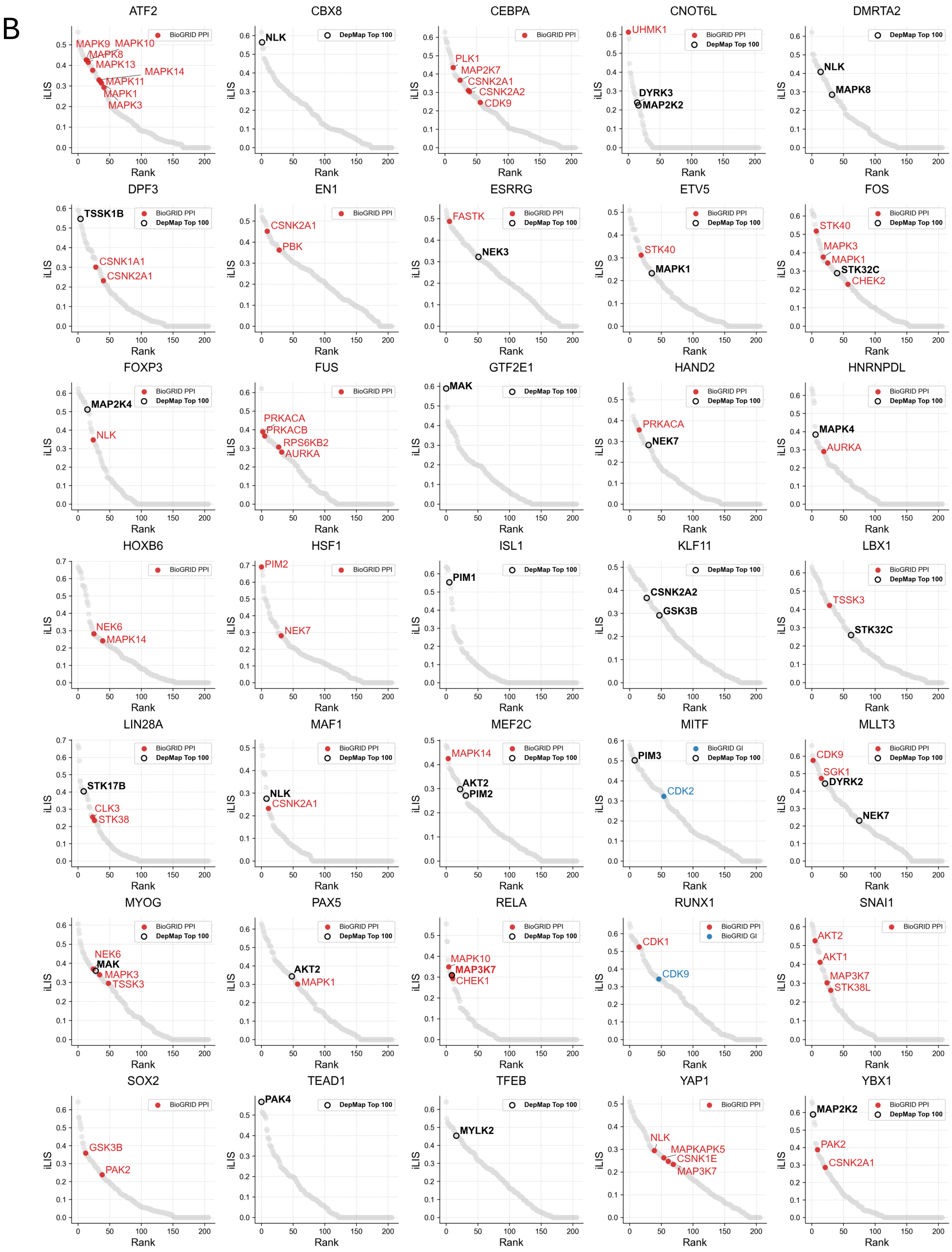

### SF 5

#### Supplementary Figure 5

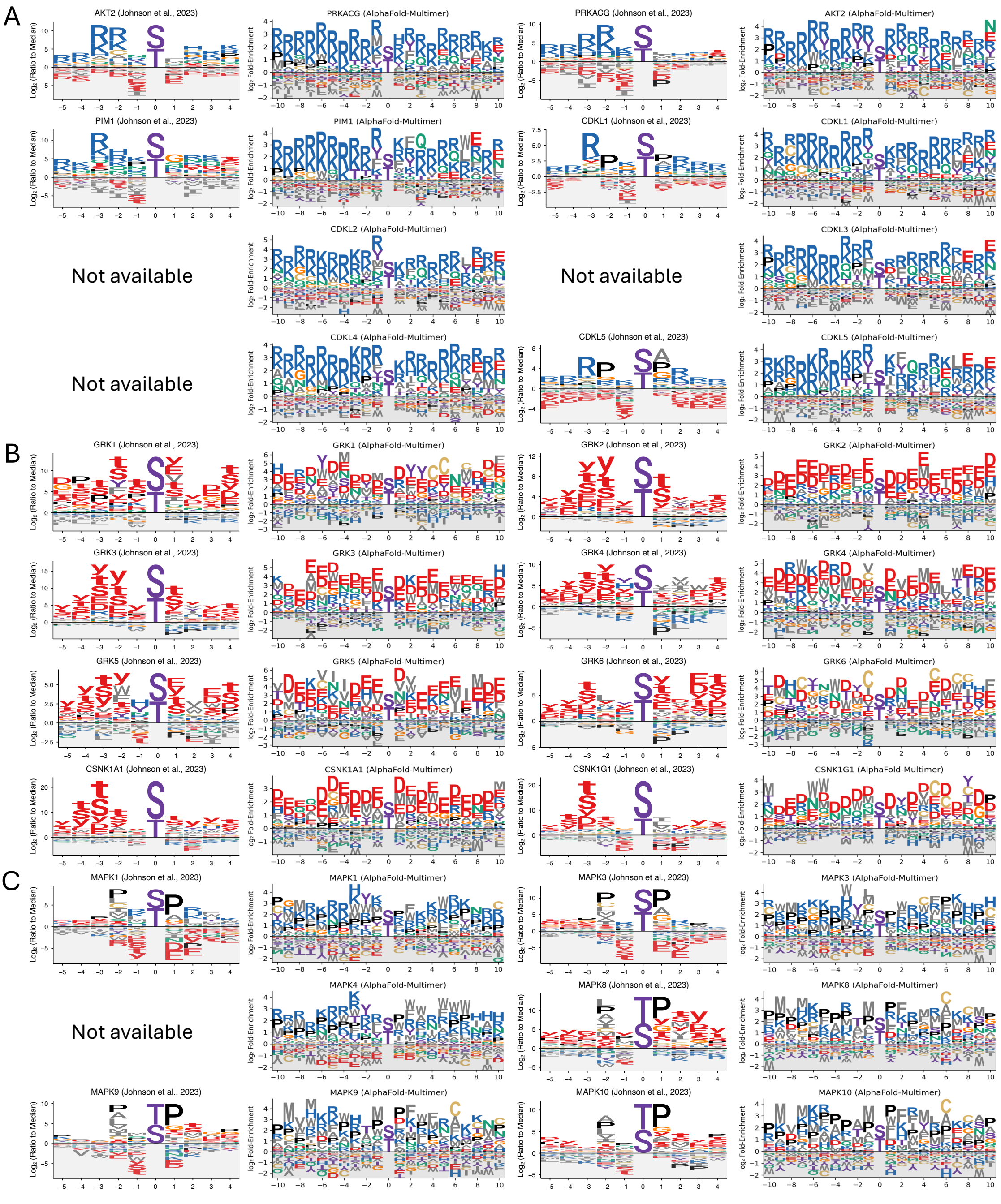

### SF 6

#### Supplementary Figure 6

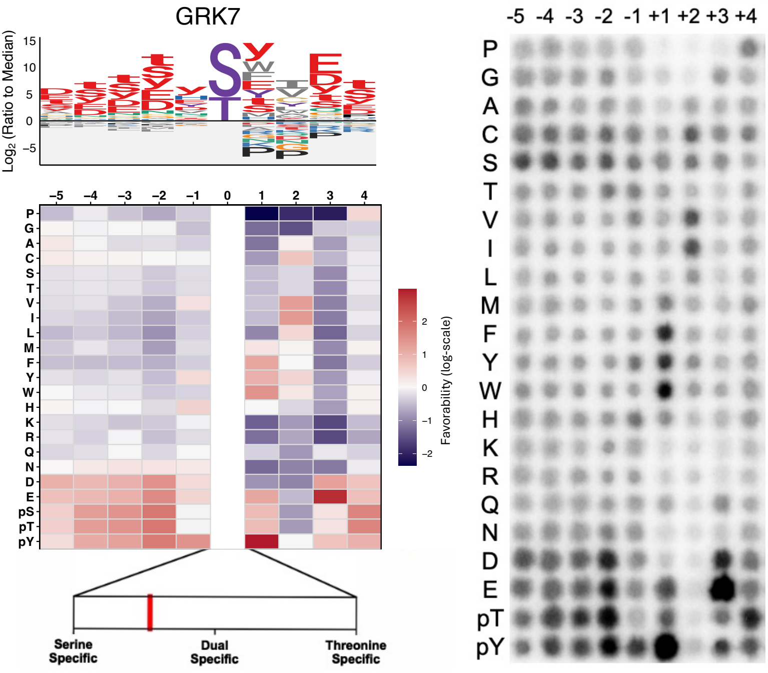

### SF 7

#### Supplementary Figure 7

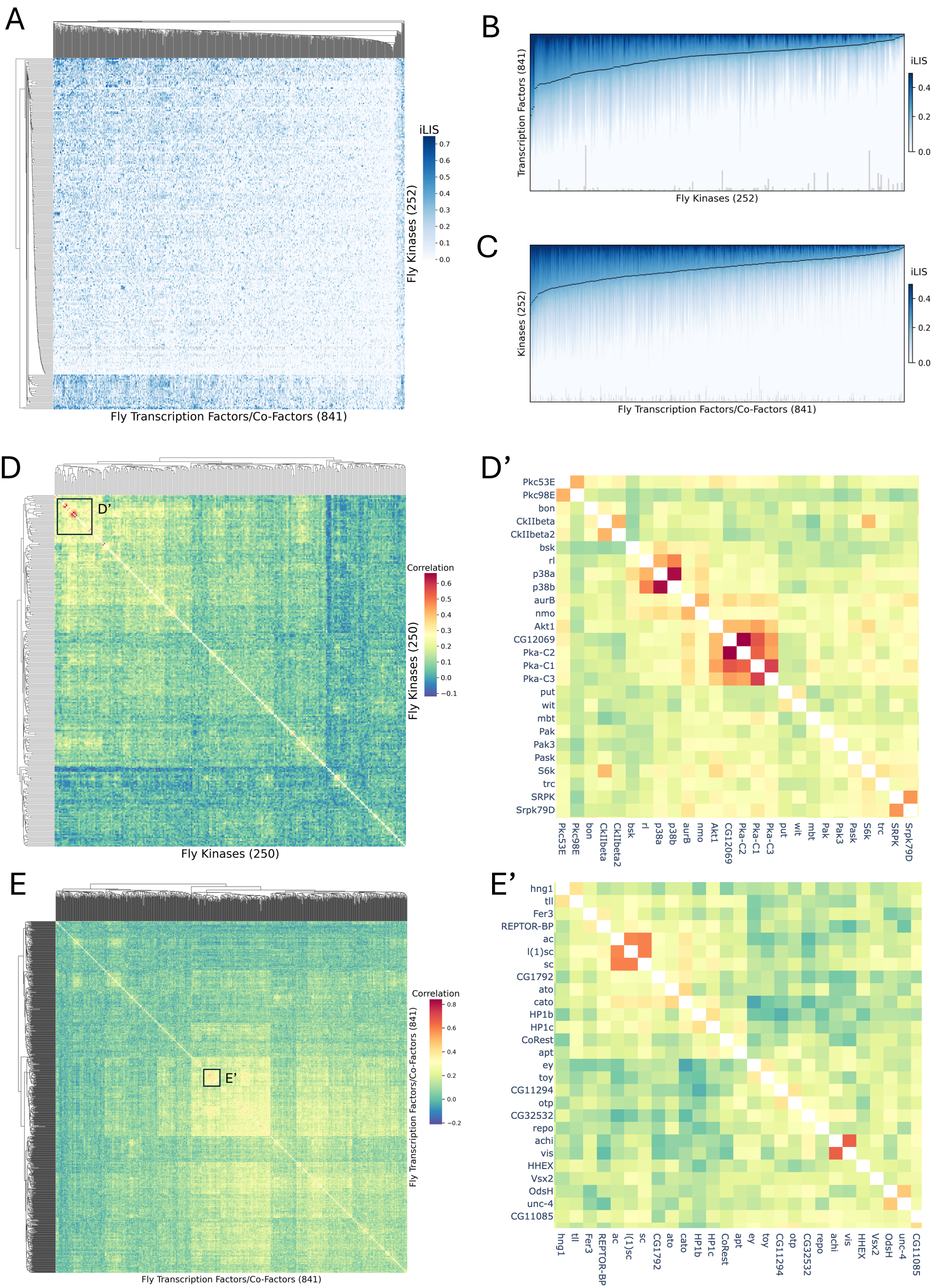

### SF 9

#### Supplementary Figure 9

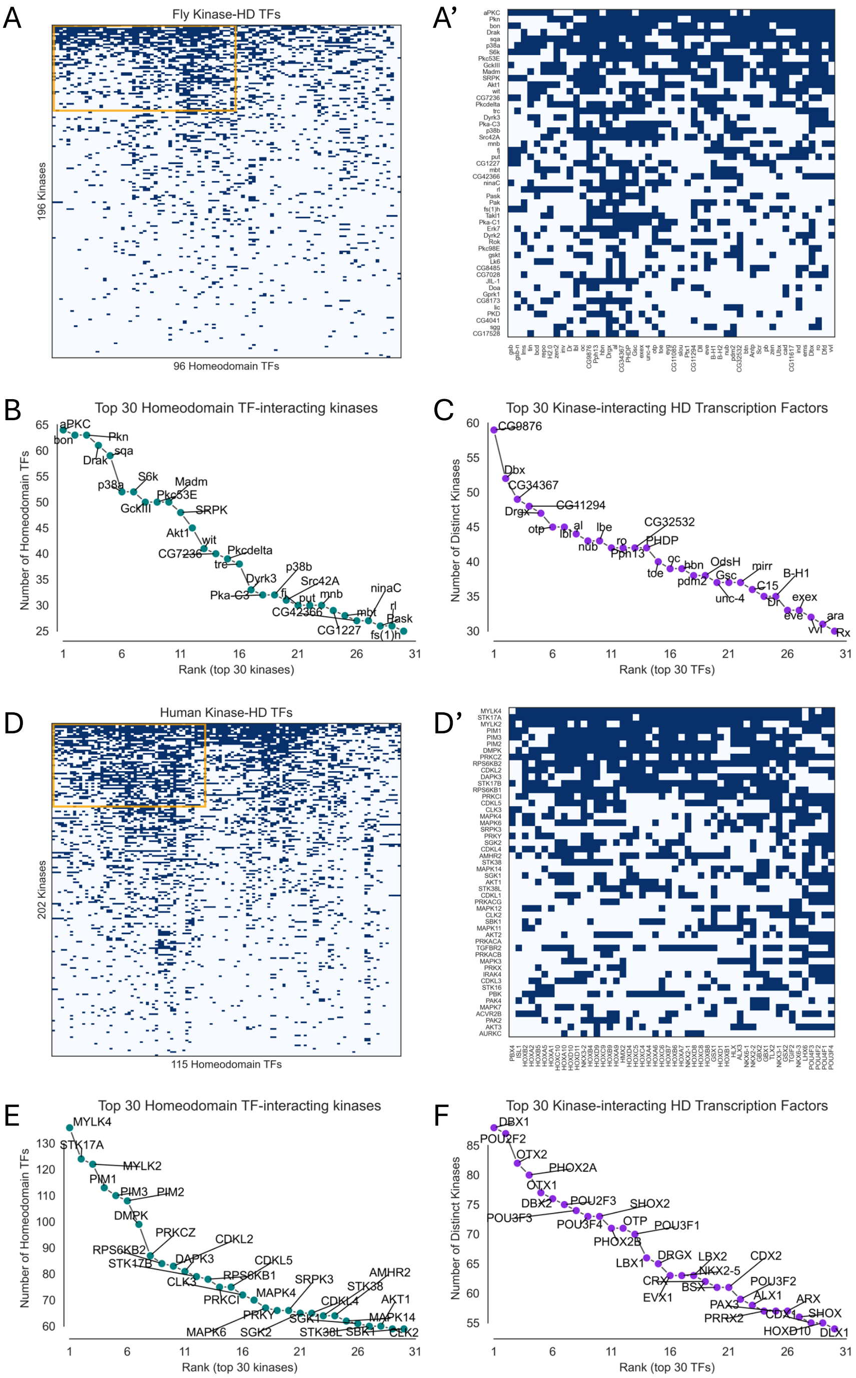

### SF 10

#### Supplementary Figure 10

A

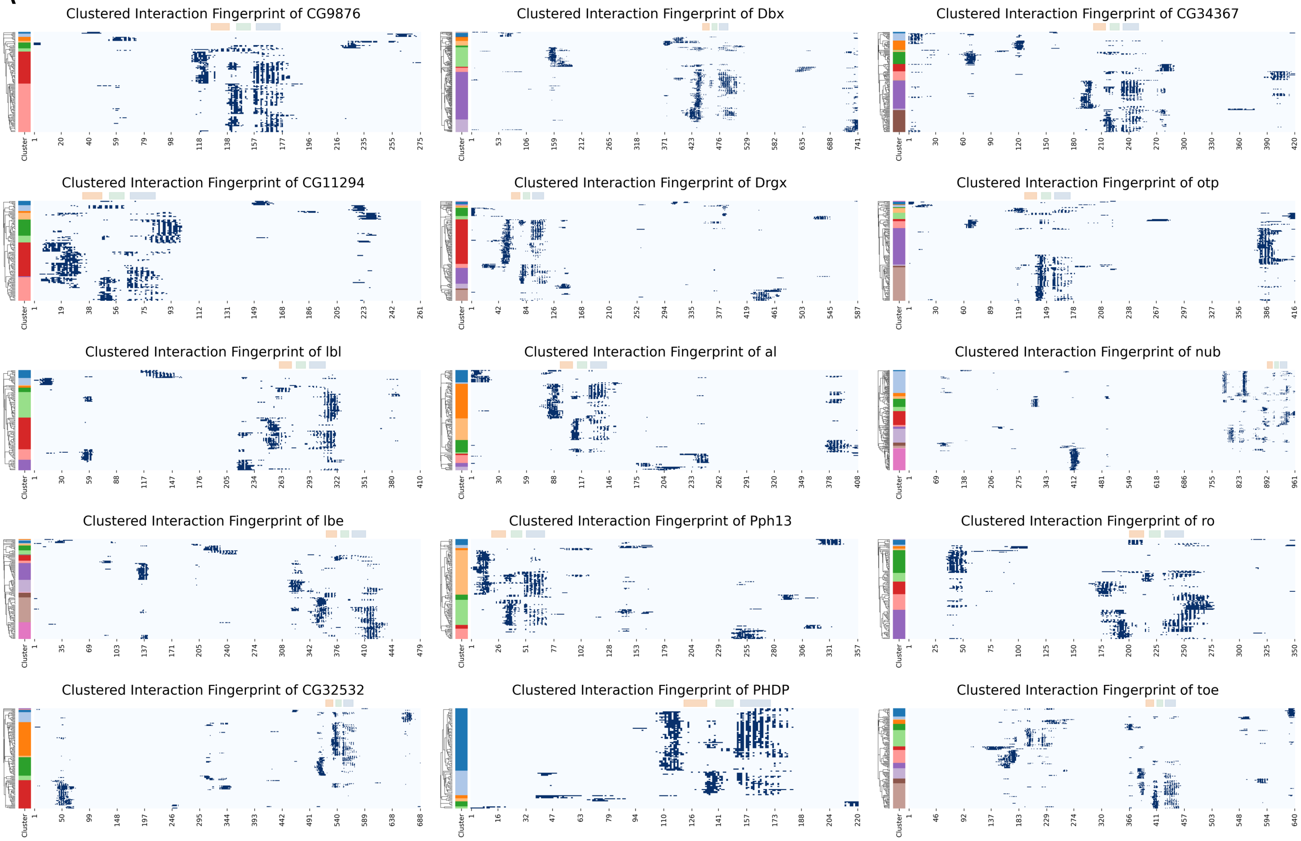


B

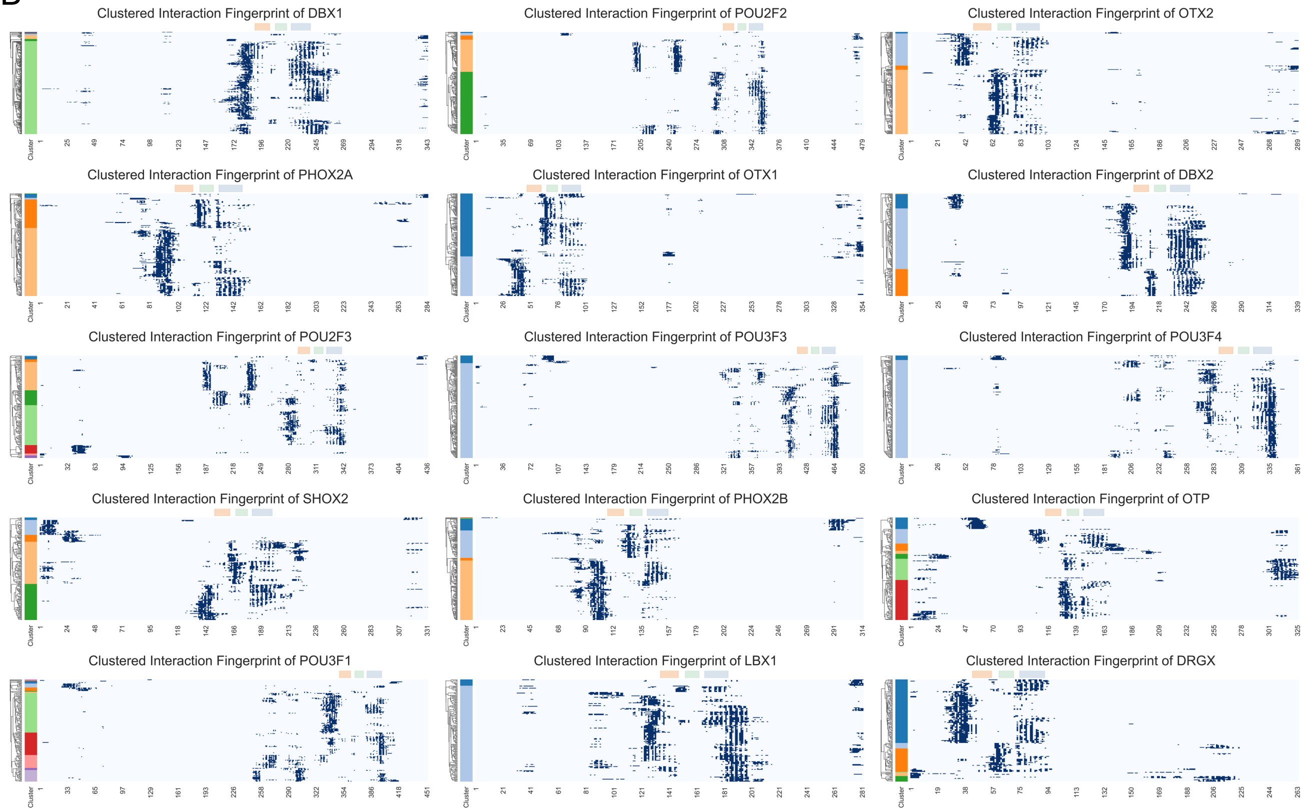

### SF 11

##### Supplementary Figure 11

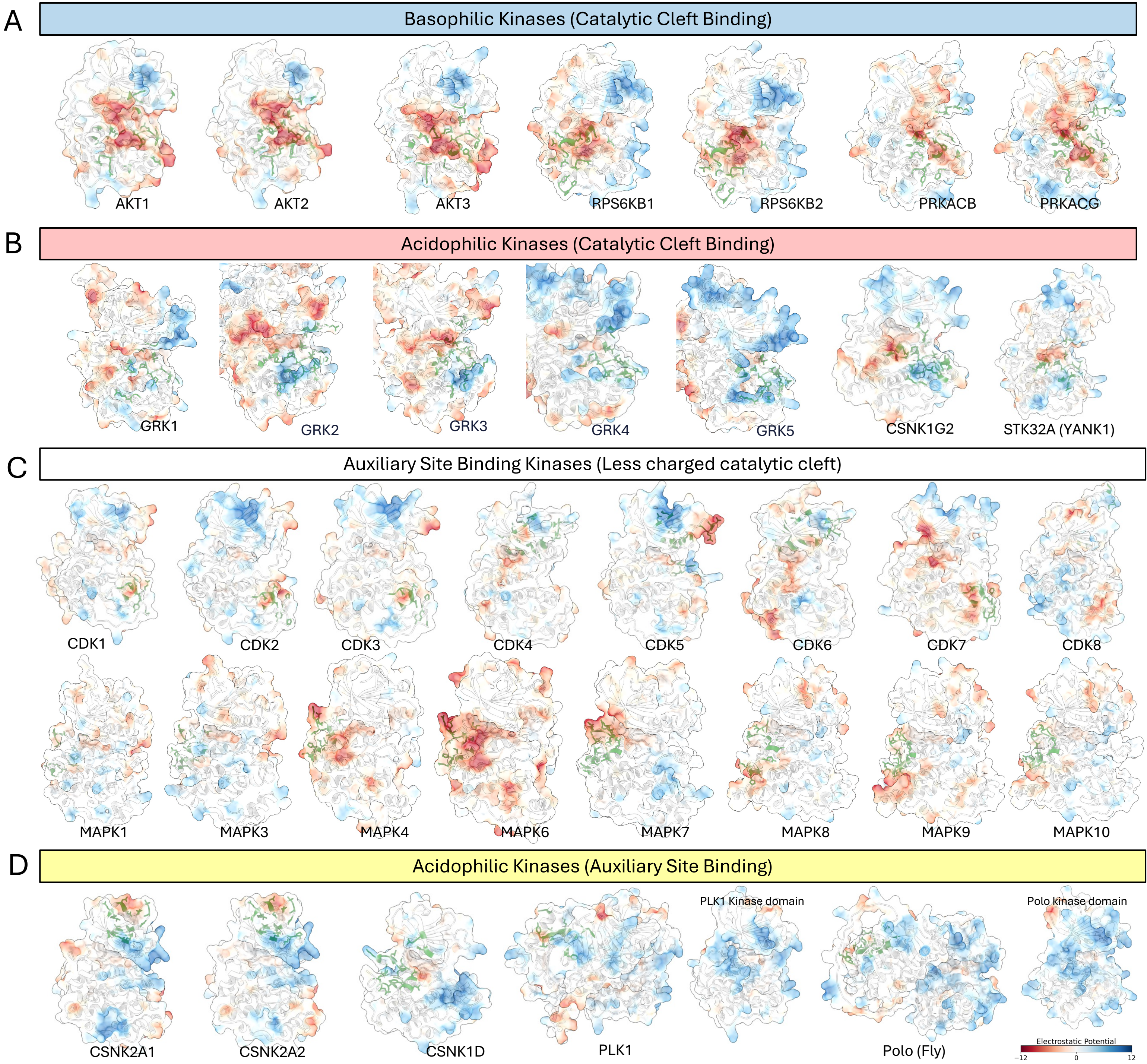

### SF 12

Supplementary Figure 12

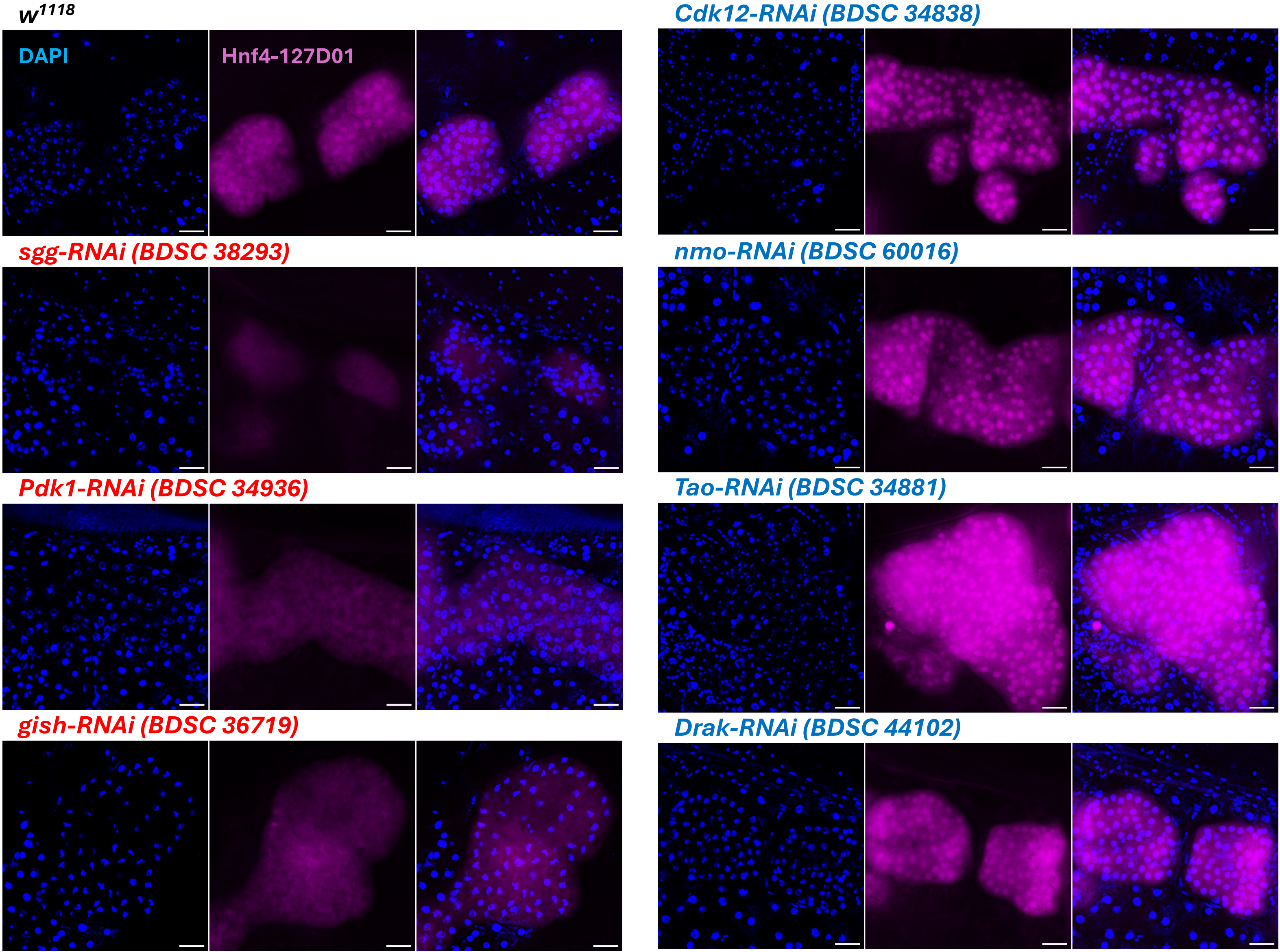
