## Supplementary material for "A Structure-Guided Kinase–Transcription Factor Interactome Atlas Reveals Docking Landscapes of the Kinome": SF 2

### Supplementary Figure 2

#### Literature-based PPIs used as Positive Reference Set in yeast Y2H screen (Yu et al. 2008)

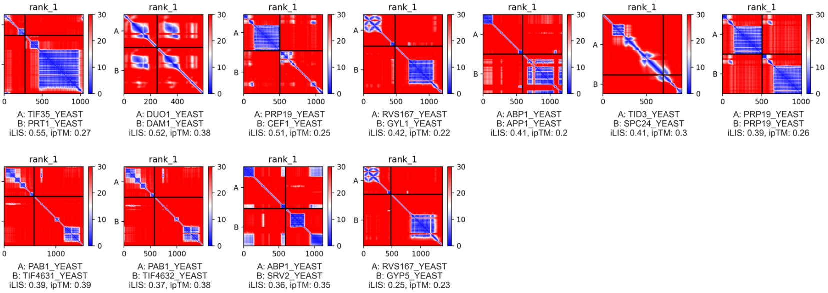


#### Literature-based PPIs used as Positive Reference Set in fly Y2H screen (Tang et al. 2023)

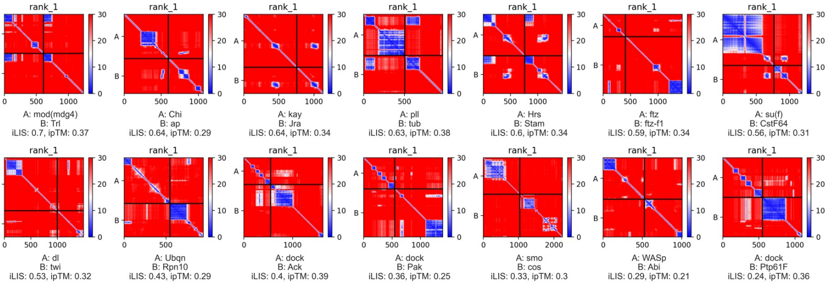


#### Literature-based PPIs used as Positive Reference Set in human Y2H screen (Braun et al. 2009)

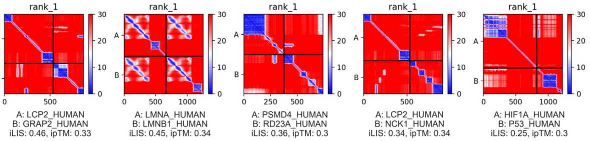


#### Structure-based PPIs annotated in Eukaryotic Linear Motif database (Kumar et al. 2024)

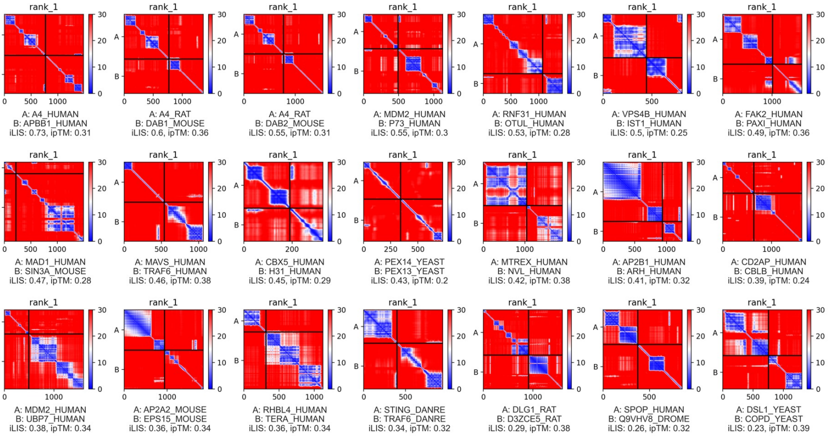
