## Supplementary material for "A Structure-Guided Kinase–Transcription Factor Interactome Atlas Reveals Docking Landscapes of the Kinome": SF 8_cluster 1

Supplementary Figure 8

Cluster 1 (14 members):
CG6800, Cdk1, Cdk12, Cdk2, Cdk4, Cdk5, Cdk7, Cdk8, Cdk9, Eip63E, Pitslre, Sdr, cdc2rk, msn

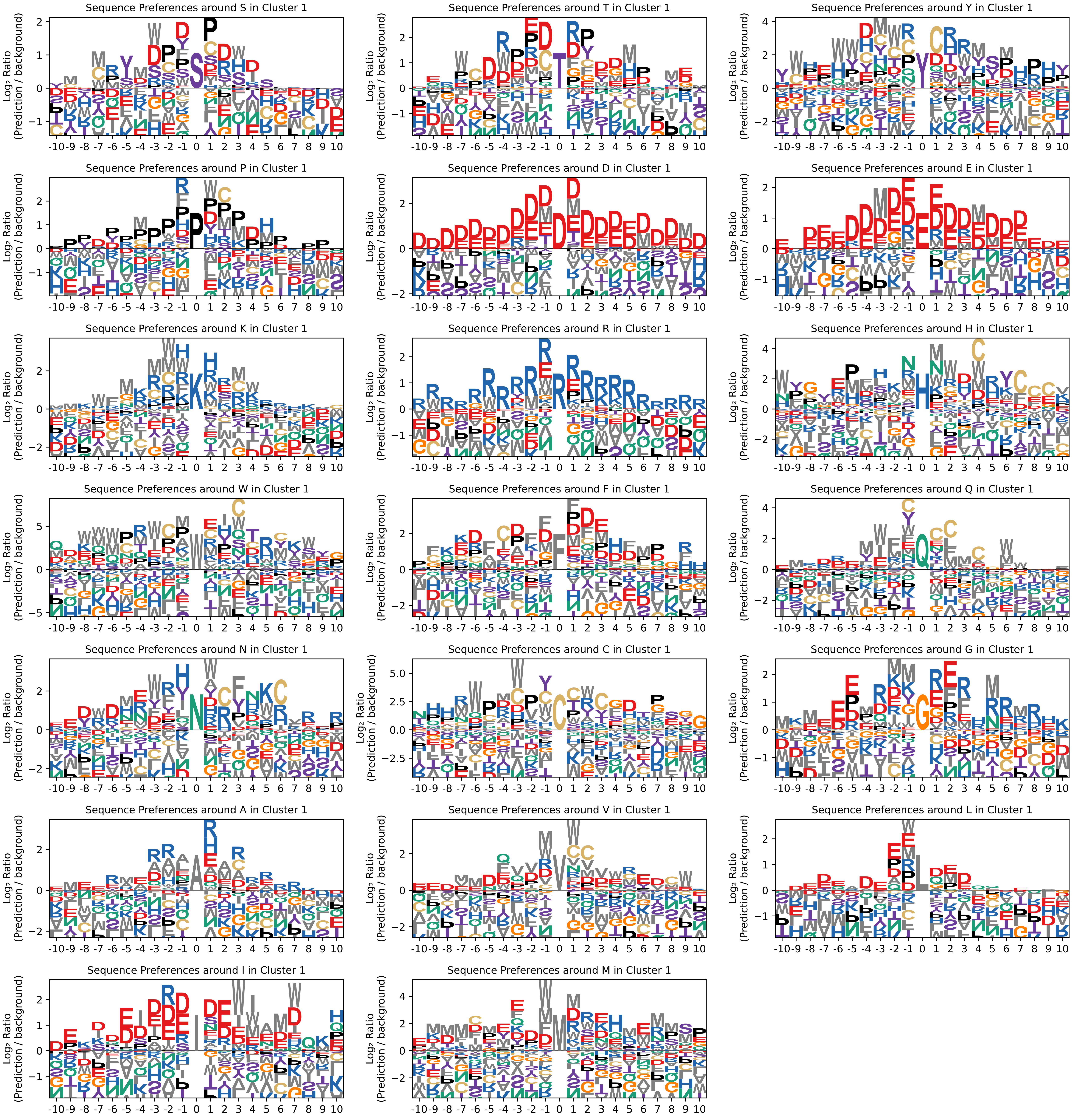
