## Supplementary figure and text_legend for "A Structure-Guided Kinase–Transcription Factor Interactome Atlas Reveals Docking Landscapes of the Kinome"

**Supplementary Text**

**1. Motivation and conceptual overview**

A primary challenge in applying AlphaFold-Multimer to signaling networks is that its standard confidence metrics, the interface-predicted TM-score (ipTM) and Model Confidence, excel at identifying rigid, stable protein complexes but systematically underestimate interactions driven by transient docking events. These include interactions mediated by short linear motifs (SLiMs) or those involving intrinsically disordered regions (IDRs), which are hallmarks of kinase-substrate recognition. The ipTM score is a global metric, meaning that long, non-binding flexible linkers or disordered tails—common in transcription factors and kinase activation loops—can dilute the confidence score, causing it to fall below the accepted threshold for a "confident" prediction (ipTM ≥ 0.7). This occurs even when the core binding interface itself is modeled with high accuracy.

To overcome these limitations of global metrics, our initial approach was to develop the Local Interaction Score (LIS), a metric that focuses only on the confidently predicted interface defined by a Predicted Aligned Error (PAE) cutoff (Kim et al., 2024). However, when applying LIS at a proteome-wide scale, we discovered a rare but significant failure mode that could lead to false positives. We observed that AlphaFold-Multimer prediction via ColabFold could occasionally predict a very low PAE for a small number of residue pairs that were physically distant and not in contact. This would result in an artificially high LIS or low minimum PAE (often used for determining positive PPIs), despite the lack of a true physical interface.

To address this specific issue, we developed the integrated Local Interaction Score (iLIS). This refined metric provides a more robust measure by ensuring that a high score reflects both high confidence and direct physical contact. iLIS is calculated as the geometric mean of the original LIS and a new metric, the contact-filtered LIS (cLIS), which is restricted to residues in direct physical proximity. The use of a geometric mean is critical: in the failure case of distant, low-PAE residues, the cLIS score will be zero, forcing the final iLIS score to zero and effectively eliminating this class of false positive. This final, balanced approach is designed to:

1. Isolate the relevant interface using the Predicted Aligned Error (PAE), a metric that reports confidence in the relative positions of all residue pairs.
2. Focus on atomic contacts through a distance-based filter, enhancing specificity.
3. Avoid rigid pLDDT cutoffs, which can inadvertently discard flexible but functionally critical interfaces common in signaling pathways.

**2. Derivation of interaction metrics**

Our structure-guided analysis relies on a suite of metrics derived from the AlphaFold-Multimer output, designed to quantify the size, quality, and confidence of predicted protein-protein interfaces.

***Core Interface Metrics (LIA, LIS, cLIA, cLIS, iLIS)***

- Local Interaction Area (LIA): It is a quantitative measure of the size of a confidently predicted binding interface. It is defined as the total number of interface residue pairs (one from each protein) where the Predicted Aligned Error (PAE) between the pair is less than or equal to 12 Å. A large LIA value suggests a broad and structurally well-defined interaction surface.
- Local Interaction Score (LIS): It measures the average confidence of the broad LIA. For each residue pair within the LIA, its PAE value is linearly inverted to a confidence score (i.e., a PAE of 0 Å yields a score of 1.0, and a PAE of 12 Å yields a score of 0). The LIS is the mean of these scores and reflects the overall structural reliability of the predicted binding interface.
- Contact-filtered Local Interaction Area (cLIA): To specifically quantify the core, physically touching interface, we defined the contact-filtered LIA (cLIA). The cLIA is defined as the total number of interface residue pairs that satisfy two criteria: they are in close physical proximity (Cβ–Cβ distance ≤ 8 Å) and their relative positions are confidently predicted (PAE ≤ 12 Å). The cLIA therefore represents the number of residue pairs predicted to be in direct, high-confidence atomic contact, helping to distinguish well-packed interfaces from more diffuse ones.
- Contact-filtered Local Interaction Score (cLIS): The contact-filtered LIS (cLIS) measures the average confidence of the core interface defined by the cLIA. It is calculated as the mean of the inverted PAE scores for only those residue pairs that constitute the cLIA. The cLIS provides a specific measure of confidence in the physically interacting residues, reducing potential inflation from confidently placed but non-contacting distal residues.
- Integrated Local Interaction Score (iLIS): To combine broad interface reliability with contact specificity, we define the final iLIS as the geometric mean of LIS and cLIS:

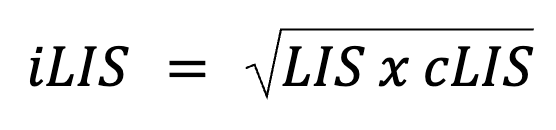

Because both LIS and cLIS range from 0 to 1, iLIS does as well. A high iLIS value indicates that an interface is predicted to be both structurally well-defined and mediated by close physical contacts, making it a robust metric for identifying high-confidence interactions.

***Interface Residue Metrics (LIR, cLIR, LIR_indice, cLIR_indice)***

To complement the pair-wise counts, we track the unique residues involved in the interaction:

- Local Interaction Residues (LIR)**:** LIR is the total count of unique residues (sum of residues from Chain A and Chain B) that participate in the LIA (PAE ≤12 Å).
- Contact-filtered Local Interaction Residues (cLIR)**:** cLIR is the total count of unique residues (sum of residues from Chain A and Chain B) that participate in the cLIA (PAE ≤12 Å AND Cβ–Cβ distance ≤8 Å).
- LIR_Indice_A and LIR_Indice_B**:** The residue indices in LIA for both interacting proteins.
- cLIR_Indice_A and cLIR_Indice_B**:** The residue indices in cLIA for both interacting proteins.

While this suite of metrics was originally designed and optimized for AlphaFold-Multimer, the underlying principles are broadly applicable. Because the framework relies on fundamental outputs like the Predicted Aligned Error (PAE) and inter-residue distances, it can be readily adapted for use with newer models such as AlphaFold 3. Moreover, the general strategy of focusing on a confidently predicted local interface, rather than relying on global scores, provides a versatile approach for interpreting predictions from other emerging structure prediction tools. It is important to note, however, that while the conceptual framework is portable, the specific cutoff values that define the method—such as the 12 Å PAE threshold, the 8 Å Cβ–Cβ distance cutoff for contacts, and the final iLIS threshold with FDR 10% (0.223)—were empirically optimized for the version of AlphaFold-Multimer used in this study. Applying this strategy to other prediction tools would necessitate a similar, rigorous benchmarking process to determine their respective optimal parameters.

**3. Optimization of the PAE threshold**

The performance of our interaction scores is critically dependent on the PAE cutoff value, C, used to define the Local Interaction Area. To determine the optimal value for C, we previously performed a systematic benchmark by calculating the Receiver Operating Characteristic Area Under the Curve (ROC AUC) for the original LIS metric across a range of PAE cutoffs from 1 Å to 30 Å. This analysis, detailed in our prior work(Kim et al., 2024), revealed a clear performance peak for both the average and best LIS metrics at a PAE cutoff of 12 Å.

Based on this established optimization, we selected 12 Å as the PAE threshold for all subsequent LIA, LIS, cLIA, cLIS, and iLIS calculations used in this study. While our comprehensive benchmark includes diverse interaction types, the specific iLIS thresholds used to classify positive PPIs were calibrated using only the large-scale Y2H reference sets (Human, *Drosophila*, and Yeast). This decision was made to tune the predictive cutoffs against datasets that most closely mimic the conditions of a large-scale discovery screen, where interactions between systematically tested, full-length proteins are most representative of a physiological search space.

**4. Benchmark dataset construction**

To rigorously train and validate our scoring method, we assembled a comprehensive benchmark dataset composed of a positive reference set and several control sets.

Positive PPI sets: Our positive set was constructed by integrating four distinct, high-quality datasets to capture a wide range of interaction types:

- Human Positive Reference Set (PRS): A set of high-quality human PPIs manually curated from literature, which originally served as a "gold standard" positive reference set to establish scoring cutoffs in a systematic yeast two-hybrid (Y2H) screen of human proteome (Braun et al., 2009).
- Fly PRS: A curated set of high-confidence *Drosophila* PPIs drawn from literature that was used as a positive control set for a large-scale Y2H screen of *Drosophila* proteome (Tang et al., 2023).
- Yeast PRS: A set of high-quality yeast PPIs from literature used to benchmark and calibrate the methodology of a large-scale Y2H study of the yeast interactome (Yu et al., 2008).
- ELM dataset: This set includes dozens of manually curated interactions from the Eukaryotic Linear Motif (ELM) database (Kumar et al., 2024). All of these interactions used in our test are validated by high-resolution experimental structures, which typically involve a truncated protein domain binding a short peptide motif. For our benchmark, we intentionally modeled the interactions using the full-length sequences of both proteins. This approach was chosen to better mimic physiological conditions and to provide a stringent test for our metric, as using the truncated sequences from crystal structures would artificially inflate structural confidence scores like ipTM.
- Time-split validation set: To assess the generalizability of our method on more recent data, we incorporated a time-split benchmark. This set consists of heterodimeric protein complexes sourced from the validation dataset of HelixFold-Multimer(Fang et al., 2024). Crucially, the experimental structures for these complexes were deposited after the training cutoff date for AlphaFold-Multimer, providing a robust test of performance on structures not seen during model training.

Control PPI sets: To assess specificity, we generated four distinct control sets:

- Random reference sets (RRS): These sets consist of protein pairs that were tested but did not show an interaction in the three large-scale Y2H screens described above. Each species-specific RRS (Human RRS, Fly RRS, and Yeast RRS) was derived from its corresponding study and served as a negative control for that screen, as the pairs were not found to interact at the time of curation (Braun et al., 2009, Tang et al., 2023, Yu et al., 2008).
- Shuffled reference set (SRS): This set was generated by randomly shuffling the protein pairs from within each of the positive datasets.
- GFP control set: Each protein from the PRS was paired with Green Fluorescent Protein (GFP). As a non-endogenous protein, GFP serves as a stringent control against non-specific binding.
- Subcellular compartment control set: Each intracellular protein from the PRS was paired with Wingless (Wg, fly Wnt), a secreted signaling protein. As these proteins reside in different cellular compartments, they provide a biologically relevant control.
- Time-split control set: Each protein from the positive Time-Split Validation Set was paired with putatively non-interacting partners, GFP and Wg to generate a corresponding time-split control set for assessing generalizability.

**5. Limitations of the benchmark dataset**

While our benchmark sets were constructed using established, high-quality sources, we acknowledge several inherent limitations. These caveats should be considered when interpreting the performance of our method.

Positive reference set (PRS): Although the PRS is built from interactions with multiple layers of evidence, not all pairs are supported by direct structural data. Consequently, for some interactions, it remains uncertain whether they are direct binary contacts or indirect associations within a larger complex. Our method is designed to predict direct interactions, which could lead to apparent false negatives for pairs that are part of the same functional complex but do not physically touch.

Control PPI sets (RRS and SRS): The "negative" classification of the RRS is based on the absence of a detected interaction in specific high-throughput screens at the time of their curation. Similarly, the SRS is based on random, operating under the assumption that a randomly chosen pair is unlikely to be a specific biological interactor. It is possible that some of these pairs are, in fact, true biological interactions that were either missed by the original screens or have yet to be discovered. The absence of evidence is not definitive evidence of absence.

Exogenous and compartmental controls: We assume that GFP and Wg do not form specific, high-affinity interactions with the intracellular proteins in our PRS. However, it is known that even standard control proteins like GFP can sometimes be "sticky" or engage in unexpected, low-affinity interactions. Furthermore, the GFP construct used in our study included N-terminal (127D01-tag) and C-terminal (VHH05-tag) nanotags (Xu et al., 2022). Our assumption was that none of the three components (GFP, N-tag, or C-tag) would interact with the PRS proteins, but this cannot be fully guaranteed.

**6. Derivation of interaction motifs from structural models**

*6.1 Rationale and overview*

A central goal of our structure-guided atlas is to move beyond binary interaction mapping to reveal the specific residue-level patterns that govern kinase recognition. To this end, we developed a computational pipeline to derive interaction motifs directly from our AlphaFold-Multimer models. This approach was applied in two main contexts. First, as a validation of our structure-guided method, we generated motifs for the human kinase–TF screen and found that these *in silico*-derived motifs were largely consistent with known, experimentally determined consensus sequences. Second, we deployed this pipeline on our comprehensive *Drosophila* kinome atlas to help define substrate specificities and uncover novel docking motifs at a systems-level scale.

*6.2 In silico identification of putative kinase substrates and binding partners*

Many kinase-substrate interactions are characterized by the binding of a structured kinase domain to a flexible region on its partner. To computationally enrich this class of interaction, we used a filtering strategy based on the local confidence of the predicted interface residues. We use the contact-filtered Local Interaction pLDDT (cLIpLDDT), which is the average confidence score of only those residues in direct physical contact at the predicted interface. For each kinase-partner complex, the partner was classified as a putative substrate if it met the following criterion:

cLIpLDDT(Kinase) − cLIpLDDT(Partner) ≥ 15

This criterion is designed to enrich for interactions with a significant confidence differential at the binding interface, a structural signature consistent with kinase-substrate recognition. All protein pairs passing this filter were carried forward for motif analysis.

*6.3 Construction of proteome-wide background frequency models*

The raw frequency of amino acids observed in a set of binding partners can be biased by the natural abundance of those amino acids. To distinguish sequence preference from this background noise, we generated comprehensive background models. This procedure was performed independently for both the human and *Drosophila* proteomes. The human background model was built using the complete set of reviewed protein sequences from UniProt, while the *Drosophila* model was built using a comprehensive set of longest-isoform protein sequences. For each proteome, the algorithm located every instance of each of the 20 standard amino acids and tabulated the frequencies of its neighboring residues, resulting in a position-specific background model that captures the expected occurrence of any amino acid surrounding a given central residue.

*6.4 Calculation of background-corrected position weight matrices (PWMs)*

With a set of putative binding partners for each kinase and a corresponding proteome-wide background model, we calculated the final PWMs.

- Predicted frequencies: For a given kinase, we aggregated all of its identified binding partners. The interaction sites on each partner were defined by the set of contact-filtered Local Interaction Residues (cLIRs). To create a contiguous motif region from potentially fragmented contacts, we implemented a gap-filling procedure: if the sequence distance between two cLIRs was five or fewer residues, the intervening residues were also included in the site definition. After defining these contiguous interaction sites, we located every potential central residue in a departure from conventional phosphosite-centric analysis, performing our analysis systematically by centering on each of the 20 standard amino acids. We then collected the flanking sequences (±10 residues) to generate an "predicted" frequency matrix.
- Fold-enrichment normalization: To correct for background abundance, the predicted frequency of each amino acid at each position was divided by its corresponding background frequency:

*Fold-Enrichment = Predicted Frequency / Background Frequency*

To avoid division-by-zero errors, a small epsilon value was used to impute any zeros in the background model. The final matrix was then converted to a log₂ scale, where positive values represent relative enrichment and negative values represent relative depletion.

*6.5 Visualization and motif averaging*

To visualize the final motifs, we applied several quality-control and presentation steps.

- Quality control: PWMs were generated only if the underlying motif was derived from a minimum of 15 unique peptide sequences to increase the robustness of the result.
- Motif averaging: For canonical phosphosite motifs for serine/threonine kinases, the individual log₂-enrichment PWMs for different acceptor types (serine and threonine) were averaged to produce a single, representative consensus motif.
- Logo generation: Sequence logos were created using the logomaker library(Tareen and Kinney, 2020) with several custom visualizations. An artificial spike was inserted at position 0 to mark the central residue of the motif, and the y-axis was scaled asymmetrically to visually emphasize residues that are positively selected.
- The complete set of derived motifs for both the human and *Drosophila* kinomes, centered on all 20 amino acids, is provided as a comprehensive resource in the Supplementary Data 2 and 4.

*6.6 Limitations of the motif derivation method*

It is important to acknowledge the inherent limitations of this computational approach.

- The cLIpLDDT heuristic: The cLIpLDDT >= 15 criterion is a heuristic designed to enrich for a specific interaction topology (a stable domain binding a flexible partner). It does not guarantee a functional kinase-substrate relationship, as some partners that pass this filter may be non-catalytic interactors, pseudosubstrates, or inhibitors. Conversely, this filter will likely exclude genuine interactions that occur between two well-structured and rigid proteins.
- Assumption of binary interactions: Our method models direct, binary interactions in isolation. It cannot account for interactions that depend on cellular context, such as those mediated by a third-party scaffold or bridging protein.
- Absence of post-translational modifications (PTMs): The structural models were generated using unmodified protein sequences. The analysis, therefore, does not account for the role of PTMs, such as priming phosphorylations, which are critical for substrate recognition by some kinase families (e.g., GSK3, Casein Kinases). The derived motifs represent the intrinsic specificity for unmodified sequences.

**7. Unsupervised clustering and structural interactome tree construction**

To classify kinases based on their substrate interaction patterns, we developed a sophisticated, multi-stage unsupervised clustering pipeline. This approach was designed to be robust to parameter choices and ensures the resulting functional classification is based on stable, reproducible features in the data.

*Step 1: Data filtering and fingerprint matrix construction*

To create a high-quality dataset suitable for our computationally intensive clustering pipeline, we first addressed the variable data coverage inherent in a screen of this scale. We established a minimum data coverage threshold, retaining only those kinases for which at least 300 TF interaction models were generated and those TFs for which at least 200 kinase interaction models were generated. This filtering process retained 231 kinases for analysis, including pseudokinases.

From this high-coverage set, we constructed a high-dimensional "TF-binding fingerprint" for each kinase. The features of this fingerprint are the complete set of unique TF residues that were identified as contact-filtered Local Interaction Residues (cLIRs) in at least one model. To create a more robust fingerprint that rewards consistently predicted contacts, we integrated information across all five generated models for each kinase-TF pair, rather than relying on a single top-ranked structure. The value for each feature (kinase_i, TF_residue_j) was calculated as the sum of the iLIS scores from every model in which that specific residue was identified as a cLIR (PAE ≤ 12 Å and Cβ-Cβ distance ≤ 8 Å), as formalized by the following equation:

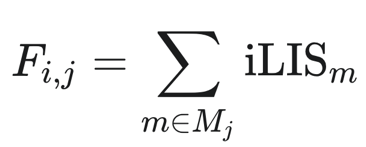

Where:

- *F_i,j_* is the final feature value for the interaction between kinase *i* and TF residue *j*.
- *m* is an individual model from the set of N models generated (where *N*=5).
- *iLIS_m_* is the iLIS score of a specific model *m*.
- *M_j_* is the set of all models where residue *j* meets the cLIR criteria.

This summation approach gives a higher weight to residues that are repeatedly predicted to be part of the binding interface. This process resulted in a large, sparse input matrix of 231 kinases × 219,796 features, which was then filtered one last time to ensure all kinases were present in our final curated annotation list.

*Step 2: Generation of a consensus co-occurrence matrix via stability analysis*

To derive a stable clustering solution that is insensitive to the stochastic nature of dimensionality reduction algorithms and specific hyperparameter choices, we employed a large-scale consensus approach. We executed a core analysis pipeline a total of 10,800 times, aggregating the results to measure the functional similarity between kinases.

The pipeline consisted of three stages with systematically varied hyperparameters. First, the high-dimensional fingerprint matrix was projected into a 2D embedding using the UMAP algorithm(McInnes et al., 2020) with a cosine similarity metric; across the stability analysis, we varied UMAP's min_dist (0.01, 0.05, 0.1, 0.15) and n_neighbors (5, 10, 15). Second, clusters were identified within the resulting UMAP embedding using the HDBSCAN algorithm(McInnes et al., 2017), varying the min_samples parameter (2, 3, 5). Finally, to ensure every kinase was assigned to a functional group, any points classified as noise by HDBSCAN were rescued by assigning them to the majority cluster of their k-nearest neighbors, with k also varied (1, 2, 3) for the rescue.

*Step 3: Aggregation into a consensus matrix*

The clustering assignments from all 10,800 runs were aggregated into a single, final consensus co-occurrence matrix. Each cell in this matrix stores the fraction of runs in which two kinases were assigned to the same cluster, providing a highly robust, parameter-agnostic measure of their functional similarity. The stability of the clustering across the parameter space was assessed by calculating the Adjusted Rand Index (ARI) for each bootstrap replicate against the result from the full dataset.

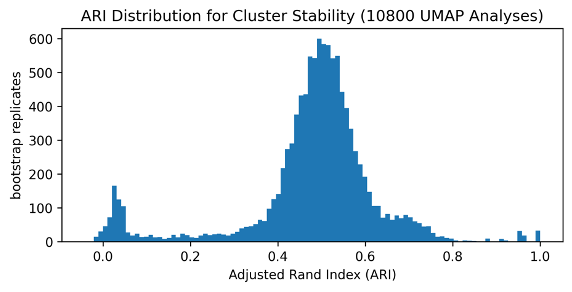

*Step 4: Hierarchical clustering and final tree generation*

The final dendrogram and cluster assignments were generated from the consensus co-occurrence matrix using a multi-step process in R.

- Distance calculation and hierarchical clustering: The co-occurrence matrix was first normalized by its maximum value. It was then converted to a distance matrix using the formula distance = 1 - normalized_similarity. This distance matrix served as the input for agglomerative hierarchical clustering using a complete-linkage method (hclust function, method="complete").
- Dynamic tree cutting: Rather than cutting the tree at a fixed number of clusters, we used a more sophisticated, data-driven approach to define the final kinase families. We applied the Dynamic Tree Cutting algorithm (R package cutreeDynamic)(Langfelder et al., 2008) to the dendrogram. This adaptive method identifies "natural" clusters by analyzing the shape of the dendrogram's branches, without requiring a pre-specified number of clusters (k). The algorithm was configured with a minimum cluster size of 5 and a deepSplit parameter of 0 to favor larger, more distinct clusters.
- Cluster relabeling and visualization: The clusters identified by cutreeDynamic were systematically re-labeled for clear and intuitive visualization in the final circular dendrogram. The mean angular position of all members of a given cluster was calculated, and the clusters were then renumbered sequentially based on their clockwise position around the tree. This ensures that adjacent wedges in the plot have consecutive numbers. The final tree was rendered using the ggtree package(Yu et al., 2017), with kinase families color-coded and pseudokinases marked with an asterisk for comprehensive annotation.

**References**

Fang, X., Gao, J., Hu, J., Liu, L., Xue, Y., Zhang, X., Zhu, K., 2024. HelixFold-Multimer: Elevating Protein Complex Structure Prediction to New Heights. https://doi.org/10.48550/arXiv.2404.10260

Kim, A.-R., Hu, Y., Comjean, A., Rodiger, J., Mohr, S.E., Perrimon, N., 2024. Enhanced Protein-Protein Interaction Discovery via AlphaFold-Multimer. https://doi.org/10.1101/2024.02.19.580970

Langfelder, P., Zhang, B., Horvath, S., 2008. Defining clusters from a hierarchical cluster tree: the Dynamic Tree Cut package for R. Bioinformatics 24, 719–720. https://doi.org/10.1093/bioinformatics/btm563

McInnes, L., Healy, J., Astels, S., 2017. hdbscan: Hierarchical density based clustering. Journal of Open Source Software 2, 205. https://doi.org/10.21105/joss.00205

McInnes, L., Healy, J., Melville, J., 2020. UMAP: Uniform Manifold Approximation and Projection for Dimension Reduction. https://doi.org/10.48550/arXiv.1802.03426

Tareen, A., Kinney, J.B., 2020. Logomaker: beautiful sequence logos in Python. Bioinformatics 36, 2272–2274. https://doi.org/10.1093/bioinformatics/btz921

Xu, J., Kim, A.-R., Cheloha, R.W., Fischer, F.A., Li, J.S.S., Feng, Y., Stoneburner, E., Binari, R., Mohr, S.E., Zirin, J., Ploegh, H.L., Perrimon, N., 2022. Protein visualization and manipulation in Drosophila through the use of epitope tags recognized by nanobodies. eLife 11, e74326. https://doi.org/10.7554/eLife.74326

Yu, G., Smith, D.K., Zhu, H., Guan, Y., Lam, T.T.-Y., 2017. ggtree: an r package for visualization and annotation of phylogenetic trees with their covariates and other associated data. Methods in Ecology and Evolution 8, 28–36. https://doi.org/10.1111/2041-210X.12628

**Supplementary Figure 1. iLIS outperforms structural confidence metrics for PPI prediction with superior performance on flexible interactions.**

(A, B) Pearson correlation matrices comparing iLIS to structural (ipTM, Model Confidence, pDockQ, pDockQ2) and recent (actifpTM, ipSAE) local confidence metrics. Scores are shown for the single best-ranked model (A) and the average of five models (B) for each interaction pair. The best-ranked model is determined by the ipTM score in ColabFold.

(C, D) Distribution of scores for each metric across the positive and control datasets, stratified by pLDDT group. The violin plots show that local confidence metrics (iLIS, actifpTM, and ipSAE) provide better separation between positive (orange) and control (blue) PPIs, particularly in the low-pLDDT (0-50) group representing flexible interactions. Scores are shown for the single best-ranked model (C) and the average of five models (D).

(E, F) Receiver Operating Characteristic (ROC) curve analyses comparing the performance of iLIS and other metrics across the full benchmark and stratified by pLDDT subgroups. Panel (E) shows the performance for the single best-ranked model, while panel (F) shows the performance for the average of five models.

(G, H) Comparison of the statistical power of each metric to distinguish between positive and control PPI sets. The bar plots show the -log10(p-value) from a Mann-Whitney U test for each metric, stratified by pLDDT group. A higher bar indicates a more significant separation. Panel (H) shows the analysis for the single best-ranked model, and panel (I) shows the analysis for the average of five models.

(I, J) Bar plots comparing the True Positive Rate (TPR) at low False Positive Rates (FPR) of 10% across pLDDT subgroups for the best-ranked model (J) and the average of five models (K).

(K) A joint plot visualizing iLIS versus ipTM scores on the time-split benchmark. The central scatter plot shows each protein-protein interaction, colored by its classification as a known positive interaction (blue) or a negative control (red). For the benchmark, negative pairs were generated by pairing proteins from the positive set with putatively non-interacting partners (GFP and Wg). The marginal axes display the distribution densities for each metric, separated for the positive and negative datasets.

(L) ROC analysis comparing the performance of iLIS versus ipTM on the low-confidence subset (pLDDT 0–50) of the time-split benchmark.

(M) Bar plots showing the True Positive Rate (recall) for each metric at high-confidence False Positive Rates (FPR) of 10% on the time-split benchmark.

**Supplementary Figure 2. iLIS rescues the prediction of transient and flexible interactions that receive low ipTM scores.**

Predicted Aligned Error (PAE) plots for a representative set of bona fide protein-protein interactions (PPIs) from the positive PPI datasets. These interactions, which are scored highly by iLIS despite low global confidence scores (ipTM), exhibit a characteristic PAE signature: a localized region of high confidence (blue area) within a globally uncertain prediction (red background). The calculated iLIS and ipTM scores are displayed below each plot. Interactions were sourced from: literature-based PPIs from Positive Reference Sets used in yeast two-hybrid (Y2H) screen for yeast (Yu et al., 2008), *Drosophila* (Tang et al., 2023), and human (Braun et al., 2009), and structurally-defined interactions from the Eukaryotic Linear Motif (ELM) database (Kumar et al., 2024).

**Supplementary Figure 3. A structure-guided interactome of the human Ser/Thr kinase–transcription factor.**

(A) A comprehensive interactome of all 240,000+ predicted human kinase-TF interactions. (A') A zoomed-in view of CDKs and Cyclins and 14-3-3 proteins. (A'') A zoomed-in view highlighting several MAPKs and FOXO family transcription factors.

(B) Kinase correlation map by transcription factor interaction profile. (B’) A zoomed-in view of MAPKs, PAKs, and S6Ks. (B’’) A zoomed-in view of CDKs.

(C) TF correlation map by kinase interaction profile. (C’) A zoomed-in view of HOX family transcription factors. (C’’) A zoomed-in view of FOXO family transcription factors.

**Supplementary Figure 4. Validation of the kinase-TF atlas against known interaction and functional co-dependency datasets.**

(A) Interaction profiles for representative human S/T kinases screened against transcription factors/co-factors. Each plot ranks partners by their iLIS score. The annotation scheme applies to both (A) and (B): Previously reported interactions in BioGRID are indicated by colored dots for Protein-Protein Interactions (PPI, red), Genetic Interactions (GI, blue), or both (purple). A black ring highlights partners that rank among the top 100 functionally co-dependent genes (ranked by absolute correlation) for that specific kinase/TF in the DepMap CRISPR screen. Highlights for external data are shown only for predictions meeting the threshold of iLIS >= 0.223.

(B) Interaction profiles for representative human transcription factors screened against human S/T kinases.

**Supplementary Figure 5. Comparison of experimental and predicted interaction motifs for basophilic, acidophilic, and proline-directed kinases.**

Side-by-side comparison of experimentally determined Position Weight Matrices (PWMs) (Johnson et al., 2023; left panels) and computationally derived PWMs from the structure-guided atlas (right panels) for additional human kinases. In the experimental PWMs, lowercase letters (s, t, y) denote the central phosphorylated Serine, Threonine, or Tyrosine residue.

(A) PWM comparisons for basophilic kinases, including members of the PKA, AKT, and CDKL families.

(B) PWM comparisons for acidophilic kinases, including members of the GRK and CSNK1 families.

(C) PWM comparisons for proline-directed kinases, including multiple members of the MAPK family.

**Supplementary Figure 6. A new experimental profiling of human GRK7 phosphosites.**

Experimental determination of the substrate specificity for the human kinase GRK7 using a peptide library array. (Top Left) The consensus motif derived from the array data is visualized as a sequence logo. (Middle Left) A heatmap shows a quantitative representation of the array data, where the favorability of each amino acid at each position is plotted on a log-scale. Red indicates high favorability (enrichment), while blue indicates low favorability (depletion). (Bottom Left) Below the heatmap, a diagram summarizes the kinase’s specificity for Serine versus Threonine at the central site. (Right) A representative image of the peptide array screen shows the raw phosphorylation data.

**Supplementary Figure 7. A structure-guided interactome of the fly kinase–transcription factor.**

(A) A comprehensive interactome of fly kinases (252) and transcription factors/co-factors (841).

(B) Accumulated iLIS distribution plot of fly kinases against transcription factors. Gray indicates no prediction due to the too large complex size.

(C) Accumulated iLIS distribution plot of fly transcription factors against kinases. Gray indicates no prediction due to the too large complex size.

(D) Kinase correlation map by transcription factor interaction profile. (D’) A zoomed-in view of kinase correlation map highlighting MAPKs (bsk, rl, p38a, p38b), AGC kinases (Akt1, Pka-C1, Pka-C2, Pka-C3).

(E) TF correlation map by kinase interaction profile. (E’) A zoomed-in view of TF correlation map showing the cluster of functionally relevant factors such as neurogenesis-related TFs (ac, l(1)sc, and sc).

**Supplementary Figure 8. The complete set of PWMs for all 20 Amino acids across all kinase clusters.**

The PWMs for the 24 binding-defined kinase clusters identified in Figure 4. Departing from a conventional focus on phosphosites, this analysis was performed systematically by centering on all amino acids (indicated above each plot). The y-axis of each logo represents the log₂ fold-enrichment for each amino acid at each position relative to its background frequency in the *Drosophila* proteome; positive values indicate enrichment. An artificial spike marks the central residue (position 0) for clarity. This comprehensive atlas enables the discovery of not only canonical substrate specificities around Serine, Threonine, and Tyrosine but also non-catalytic "microfeatures," such as the flanking enrichment of W, F, Q, and N residues that form composite docking motifs. Full methodological details are provided in Supplementary Text 6.

**Supplementary Figure 9. Widespread interactions between kinases and homeodomain transcription factors in fly and human**

(A) Interaction heatmap of fly kinases and homeodomain transcription factors (HD TFs). Kinases (Y-axis) are ordered by the number of protein-protein interactions (PPIs), and TFs (X-axis) are clustered by their interaction pattern. The orange box highlights the region shown in a magnified view in Panel A'.

(B) Top 30 HD TF-interacting kinases in *Drosophila*. Kinases are ranked by the number of unique HD TFs they interact with, with the most highly connected kinases on the left.

(C) Top 30 Kinase-interacting HD TFs in *Drosophila*. HD TFs are ranked by the number of unique kinases they interact with, with the most highly connected TFs on the left.

(D) Interaction heatmap of human kinases and HD TFs. Kinases (Y-axis) are ordered by the number of PPIs, and TFs (X-axis) are clustered by their interaction pattern. The orange box highlights the region shown in a magnified view in Panel D'.

(E) Top 30 HD TF-interacting kinases in human. Kinases are ranked by the number of unique HD TFs they interact with, with the most highly connected kinases on the left.

(F) Top 30 Kinase-interacting HD TFs in human. HD TFs are ranked by the number of unique kinases they interact with, with the most highly connected TFs on the left.

**Supplementary Figure 10. A conserved kinase docking interface on homeodomain transcription factors**

(A) Clustered interaction fingerprints of top 15 HD TFs in *Drosophila* from 10C. Each heatmap shows the interaction pattern of a single HD TF with a wide range of kinases (y-axis). Kinases are clustered based on their binding pattern, which reveals distinct kinase groups. The helices of the homeodomain are color-coded: helix 1 (pale orange), helix 2 (pale green), and helix 3 (pale blue).

(B) Clustered interaction fingerprints of top 15 HD TFs in human from Supplementary Figure 10F. These heatmaps show the clustered interaction patterns of top human HD TFs with kinases.

**Supplementary Figure 11. An additional catalog of kinase interaction hotspots.**

An expanded catalog of kinase interaction hotspots provides additional examples for the principal binding modes described in Figure 6. For each kinase, the electrostatic potential is mapped onto the surface (red: acidic; blue: basic), and the frequently used interaction residues from its largest binding partner cluster are highlighted in green. All kinases shown are human, except for Polo in panel (D), which is from *Drosophila*. Interaction hotspot residues and cluster information for all human S/T and *Drosophila* kinome can be found in Supplementary Data 5.

(A) Basophilic kinases (catalytic cleft binding): Additional examples of basophilic kinases that use their negatively charged (acidic) catalytic clefts as the primary interaction interface.

(B) Acidophilic kinases (catalytic cleft binding): Additional examples of acidophilic kinases that use their positively charged (basic) catalytic clefts for partner recognition.

(C) Auxiliary site binding kinases: An expanded set of kinases, including members of the CDK and MAPK families, that possess less-charged catalytic clefts and primarily engage partners at auxiliary (allosteric) docking sites.

(D) Acidophilic kinases (auxiliary site binding): Examples of a distinct class of kinases, including CSNK2A1 and PLK1, that recognize acidic partners but utilize auxiliary docking sites rather than their catalytic clefts.

**Supplementary Figure 12. Immunostaining from the *in vivo* kinase RNAi Screen.**

Representative images of *Drosophila* oenocytes from the kinase RNAi screen for Hnf4 regulators. For each RNAi knockdown, DAPI staining (blue) shows nuclei, and Hnf4-127D01 immunostaining (magenta) shows Hnf4 protein levels. *w^1118^* serves as the control. Oenocyte-specific RNAi knockdown was induced for each candidate kinase using the *PromE-GAL4; tub-GAL80^ts^*. Red text indicates a "positive regulator" (kinase knockdown leads to a reduction in Hnf4 protein levels), while blue text indicates a "negative regulator" (kinase knockdown leads to an increase in Hnf4 protein levels). Scale bars represent 20 µm.
